# The AGC protein kinase UNICORN controls planar growth by attenuating PDK1 in *Arabidopsis thaliana*

**DOI:** 10.1101/470732

**Authors:** Sebastian Scholz, Janys Pleßmann, Regina Hüttl, Katrin Wassmer, Balaji Enugutti, Kay Schneitz

**Affiliations:** Entwicklungsbiologie der Pflanzen, Wissenschaftszentrum Weihenstephan, Technische Universität München, Freising, Germany; Pflanzengenetik, Wissenschaftszentrum Weihenstephan, Technische Universität München, Freising, Germany; Gregor Mendel Institute of Molecular Plant Biology, Vienna, Austria

**Keywords:** AGC protein kinase, Arabidopsis, floral development, integument, ovule, PDK1, petal, planar growth, UNICORN

## Abstract

Tissue morphogenesis critically depends on the coordination of cellular growth patterns. In plants, many organs consist of clonally distinct cell layers, such as the epidermis, whose cells undergo divisions that are oriented along the plane of the layer. The developmental control of such planar growth is poorly understood. We have previously identified the Arabidopsis AGCVIII-class protein kinase UNICORN (UCN) as a central regulator of this process. Plants lacking *UCN* activity show spontaneous formation of ectopic multicellular protrusions in integuments and malformed petals indicating that *UCN* suppresses uncontrolled growth in those tissues. In the current model UCN regulates planar growth of integuments in part by directly repressing the putative transcription factor ABERRANT TESTA SHAPE (ATS). Here we report on the identification of *3-PHOSPHOINOSITIDE-DEPENDENT PROTEIN KINASE 1* (*PDK1*) as a novel factor involved in *UCN-*mediated growth control. PDK1 constitutes a basic component of signaling mediated by AGC protein kinases throughout eukaryotes. Arabidopsis *PDK1* is implied in stress responses and growth promotion. Here we show that loss-of-function mutations in *PDK1* suppress aberrant growth in integuments and petals of *ucn* mutants. Additional genetic, in vitro, and cell biological data support the view that UCN functions by repressing PDK1. Furthermore, our data indicate that *PDK1* is indirectly required for deregulated growth caused by *ATS* overexpression. Our findings support a model proposing that UCN suppresses ectopic growth in integuments through two independent processes: the attenuation of the protein kinase PDK1 in the cytoplasm and the repression of the transcription factor ATS in the nucleus.

**Author Summary:** Plant organs, such as petals or roots, are composites of distinct cell layers. As a rule, cells making up a layer, for example the epidermis, the outermost layer of a tissue, divide “within the plane” of the layer. This cellular behavior results in the two-dimensional sheet-like or planar growth of the cell layer. The mechanism orchestrating such a growth pattern is poorly understood. In particular, it is unclear how uncontrolled and “out-of-plane” growth is avoided. Here we provide insight into this process. Our data indicate that higher than normal activity of a central regulator of growth and stress responses results in wavy and malformed petals and in protrusion-like aberrant outgrowths in the tissue that will develop into the seed coat. It is therefore important to keep this factor in check to allow proper formation of those tissues. We further show that a protein called UNICORN attenuates the activity of this regulator thereby ensuring the sheet-like growth of young petals or the developing seed coat.

## Introduction

Spatial coordination of cell division patterns within a tissue layer is an essential feature of plant tissue morphogenesis. For example, the shoot apical meristem generates above-ground lateral organs, such as flowers, and is a composite of clonally distinct histogenic cell layers [1]. Cells of the outermost or L1 layer will contribute to the epidermis while cells of the inner L2 and L3 layers will generate the interior tissues of a lateral organ. Similarly, the Arabidopsis root consists of radial cell layers each of which originates from the activity of corresponding initial or stem cells within the root meristem [2]. The two integuments of Arabidopsis ovules constitute another example. They represent lateral determinate tissues that originate from the epidermis of the chalaza, the central region of the ovule [3, 4]. Each integument consists of a bi-layered sheet of regularly arranged cells as the cells strictly divide in an anticlinal fashion during outgrowth [3, 5, 6]. Thus, the two integuments undergo planar or laminar growth eventually surrounding the nucellus and embryo sac in a hood-like fashion. The regular cell division pattern during integument outgrowth suggests that coordinated cellular behavior across the tissue is essential for the laminar structure of the integuments.

The genetic control of planar integument growth is poorly understood [7]. Although there exists a large number of mutants with a defect in integument development, a detailed molecular and genetic framework controlling planar growth is still lacking. Interestingly, integuments of *unicorn* (*ucn*) mutants exhibit spontaneous local ectopic growth revealing a defect in the regulation of planar growth [8]. *UCN* encodes a protein kinase of the AGC VIII family [9]. Certain members of the AGC VIII family, such as D6 PROTEIN KINASE (D6PK), PINOID (PID), or WAG2, have been shown to be important for activation of polar auxin transport [10–12] raising the possibility that *UCN* mediates planar growth through the regulation of polar auxin transport. However, there is no evidence supporting this view. The available data suggest that *UCN* is not involved in polar auxin transport [9, 12–14]. Moreover, expression of *PIN-FORMED* (*PIN*) genes, encoding the classic regulators of intercellular polar auxin transport [15, 16], could not be detected during integument outgrowth [17, 18].

How does *UCN* suppress ectopic growth in integuments? So far, genetic analysis has identified *ABERRANT TESTA SHAPE* (*ATS*) as an important factor involved in the *UCN* signaling mechanism [9, 14]. *ATS* encodes a putative transcription factor that controls early integument development [19–22]. In the current model UCN controls maintenance of planar integument growth by attenuating the activity of ATS through direct phosphorylation. In the absence of wild-type UCN function, de-repression of ATS results in an altered transcriptional program that ultimately leads to ectopic local growth in integuments. ATS could potentially provide a link to auxin as there is evidence suggesting that a complex between the auxin response factor ARF3/ETTIN (ETT) and ATS controls integument initiation [22]. However, genetic data indicate that *UCN* does not control early integument initiation and functions independently of *ARF3/ETT*. Rather, the interaction between *UCN* and *ATS* is thought to be part of a later-acting mechanism that maintains planar integument outgrowth [9, 14].

*3-PHOSPHOINOSITIDE-DEPENDENT PROTEIN KINASE 1* (*PDK1*) represents another factor potentially involved in *UCN* signaling. *PDK1* encodes a basal member of the AGC protein kinase family and a master regulator of downstream AGC kinases. PDK1 is well characterized in mammalian cells where it plays a central role in connecting lipid signaling to a broad range of cellular processes [23, 24]. Amongst others, it is involved in the promotion of cell proliferation and survival [25] and is overexpressed in many different tumors [26]. Complete absence of *PDK1* function is lethal in for example fly or mouse [27, 28]. PDK1 consists of an N-terminal kinase domain and a C-terminal pleckstrin homology (PH) domain through which it binds to phospholipids. Well-studied targets include PKA, p70 ribosomal S6 kinase (S6K), or PKB/Akt. How exactly PDK1 activates its targets is substrate-specific. As a rule, PDK1 interacts with downstream AGC kinases by binding to a small hydrophobic motif (FxxF, often extended to include a phosphorylation site) termed PDK1 interacting fragment (PIF), or PIF domain, present at the C-terminus of many target AGC kinases. The PDK1 domain mediating this interaction is called PIF-binding pocket or PIF-pocket. PDK1 then activates its various substrates by phosphorylating a specific site in the activation-loop (also known as T-loop) of the target kinase domain.

In unstimulated cells animal PDK1 is found in the cytoplasm but is largely excluded from the nucleus. Upon stimulation, however, PDK1 also accumulates in the nucleus, possibly as a means to prevent activation of cytosolic signaling pathways [29–31]. Activation of PDK1 itself appears to be a dynamic process [29, 32, 33]. The current model states that PDK1 auto-phosphorylates, and is thus principally constitutively active, but is kept inactive due to auto-inhibition mediated by the PH domain. Following a stimulus primed but inactive dimers (kept together by the PH domains) become activated upon binding of the PH domain to the plasma membrane (PM). This results in stable active PDK1 monomers that phosphorylate targets, such as PKB/Akt, at the PM [33].

Despite impressive progress the function and regulation of PDK1 in plants is comparably less well understood [34–36]. Results obtained by several labs indicate that PDK1 is involved in the control of various abiotic and biotic stress responses [37–42]. Similar to the situation in animals complete removal of *PDK1* function in tomato is lethal [38]. By contrast, complete knock-out of the single *PDK1* gene in *Physcomitrella patens* does not result in lethality but causes developmental abnormalities and late growth retardation [43]. The Arabidopsis genome contains two closely homologous *PDK1* genes, *PDK1.1* and *PDK1.2* [34]. Loss-of-function *pdk1.1 pdk1.2* double mutants show only minor aberrations including somewhat stunted growth and reduced fertility. Moreover, they are impaired in growth promotion induced by the fungus *Piriformospora indica* [44]. Interestingly, a small reduction in expression of the single rice *PDK1* gene also results in moderate dwarfism [40]. These results indicate that *PDK1* is also involved in growth control and that *PDK1* requirement may vary from species to species.

PDK1 can interact with a number of AGC kinases [45], including OXI1 [40, 46], Adi3 [38] or PID [47]. In vitro data suggest that as a rule plant PDK1 activates downstream AGC kinases by binding to the target PIF domain and phosphorylating a conserved site in the activation loop [38, 41, 46, 47]. This scenario resembles the model developed for the regulation of AGC kinases by PDK1 in mammalian cells. However, this notion is likely an oversimplification as PDK1/AGC kinase interaction may be more complex and depend on the involved proteins [45–47]. Some plant AGC kinases can auto-phosphorylate in vitro and may not require PDK1 for activation (for example [9, 39, 47]). Furthermore, presence of a PIF domain does not strictly correlate with in vitro binding and activation by PDK1 or function in planta [45, 48, 49]. Taken together the available data suggest complex biochemical and biological interactions between PDK1 and AGC kinases in plants.

Here we investigated the role of Arabidopsis *PDK1* in *UCN*-dependent growth control. Our data indicate that *PDK1* is expressed in many different tissues and that PDK1 predominantly localizes to the cytoplasm. They further imply that PDK1 physically interacts with UCN in vitro and in plant cells. Genetic data suggest that *UCN* attenuates *PDK1* function in vivo and that *PDK1* is required for growth deregulation caused by overexpression of *ATS*.

## Results

### PDK1 is expressed in many tissues and localized in the cytoplasm

We first performed a basic molecular and cell biological characterization of *PDK1*. The Arabidopsis genome contains two closely homologous *PDK1* genes, *PDK1.1* (At5g04510) and *PDK1.2* (At3g10540) [34]. PDK1.1 and PDK1.2 share 91 percent identity at the amino acid level. Further sequence searches did not identify a third *PDK1*-like gene in Arabidopsis (S1 Fig).

Phenotypic analysis of T-DNA insertion lines null for each *PDK1* transcript and the corresponding double-mutants confirmed the mild phenotype of *pdk1.1 pdk1.2* double mutants reported previously [44]. Two distinctive double mutants, carrying different combinations of T-DNA insertions in *PDK1.1* and *PDK1.2*, showed only mild reduction in plant height and slightly reduced silique length (Fig 1).

**Fig. 1.**
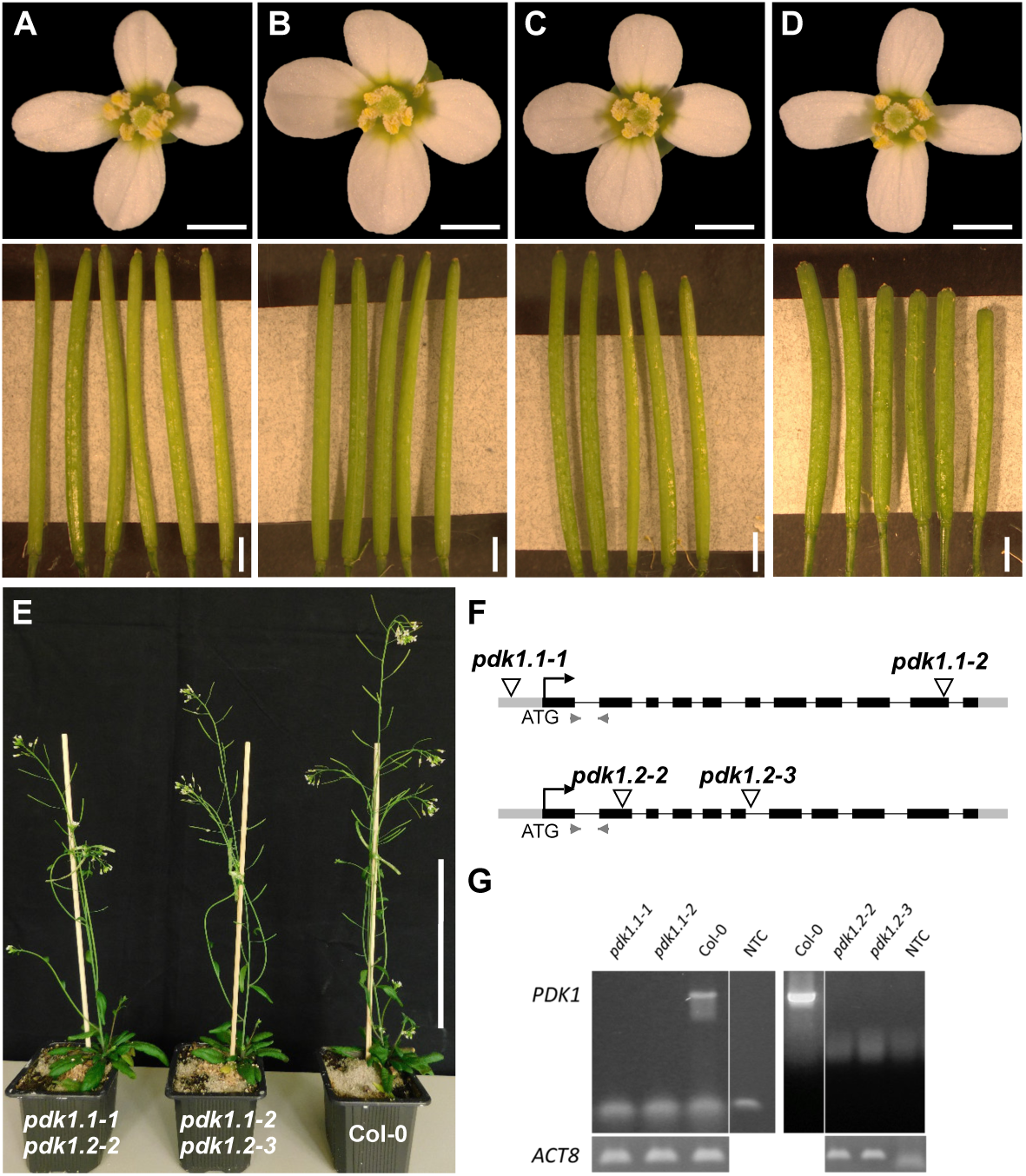
Characterization of *pdk1* T-DNA lines. Floral shapes and silique sizes of (A) Col-0, (B) *pdk1.1-1*, (C) *pdk1.2-2*, and (D) *pdk1.1-1 pdk1.2-2* T-DNA lines are shown. Note that flowers do not show an obvious aberrant phenotype. Siliques of the double mutants are shorter, slightly thicker and contain less ovules compared to WT and single insertion lines. (E) 35 days old plants of *pdk1.1-1 pdk1.2-2*, *pdk1.1-2 pdk1.2-3* and Col-0 (from left to right). Note the somewhat reduced height of the double mutants. (F) Schematic representation of the gene structure of *PDK1.1* and *PDK1.2*. Insertion sites are indicated. Grey boxes represent UTRs, black boxes exons, lines introns, and light grey arrowheads indicate primer pairs to determine transcripts in Col-0 and T-DNA lines. (G) Semi-quantitative PCR showing absence of *PDK1.1* or *PDK1.2* transcripts, respectively. Scale bars: (A-D) upper panel, 1mm; lower panel, 2 mm; (E) 10 cm.

Inspection of the AtGenExpress data set [50] revealed that both *PDK1* genes are expressed in all assayed tissues. Unfortunately, the high homology of the two *PDK1* genes rendered it impossible to perform gene-specific in situ hybridization (ISH) experiments. Nevertheless, we performed whole-mount ISH on ovules at different developmental stages. We could detect *PDK1* transcripts throughout the ovule, with the exception of the nucellus (Fig 2). We further confirmed broad expression of *PDK1.2* by assessing the tissue-level distribution of a C-terminal fusion of EGFP to PDK1.2 reporter driven by its endogenous promoter (pPDK1.2::PDK1.2:EGFP) (Fig 3). The transgene complemented the growth defects in *pdk1.1 pdk1.2* double mutants revealing its functionality (S2 Fig). We could detect expression of the pPDK1.2::PDK1.2:EGFP reporter in all assessed tissues. Moreover, we always observed reporter signal in the cytoplasm but never in the nucleus. A comparable subcellular distribution was observed using a pUBQ::PDK1.1:EGFP reporter (S3 Fig). In cells of the meristem and transition zone of roots treated with the membrane marker FM4-64 for ten minutes we also detected colocalization of PDK1:EGFP and FM4-64-derived signals (Fig 3F). Taken together, the results indicate expression of *PDK1* in all assayed tissues and a mainly cytoplasmic localization for the PDK1:EGFP fusion protein.

**Fig. 2.**
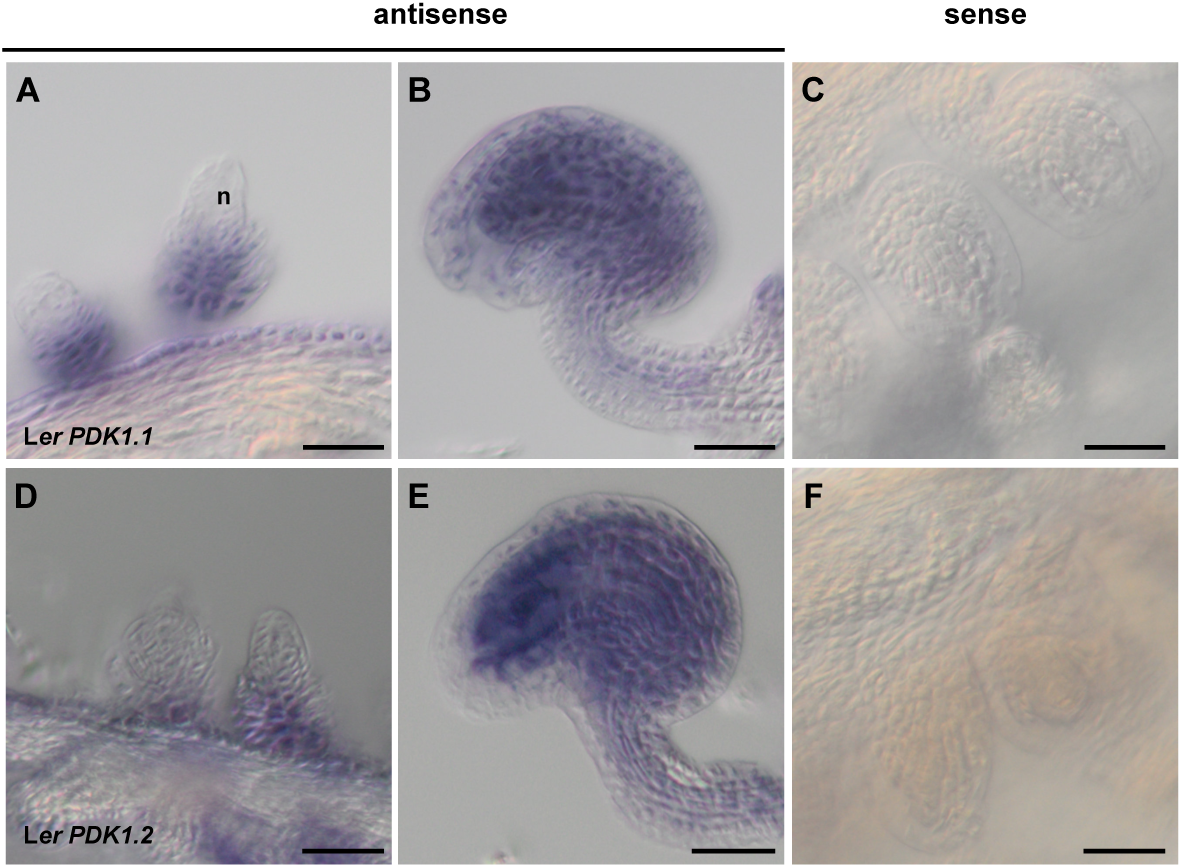
PDK1 expression in wild-type ovules detected by whole mount in situ hybridization. (A-C) *PDK1.1*. (D-F) *PDK1.2*. (A) Stage 2-III ovule (stages according to [3]). (B) Stage 3-VI ovule. (C) Stage 3-VI ovule. Sense control. Note the absence of signal. (D-F) Similar series as in (B-C). Scale bars: 25 µm.

**Fig. 3.**
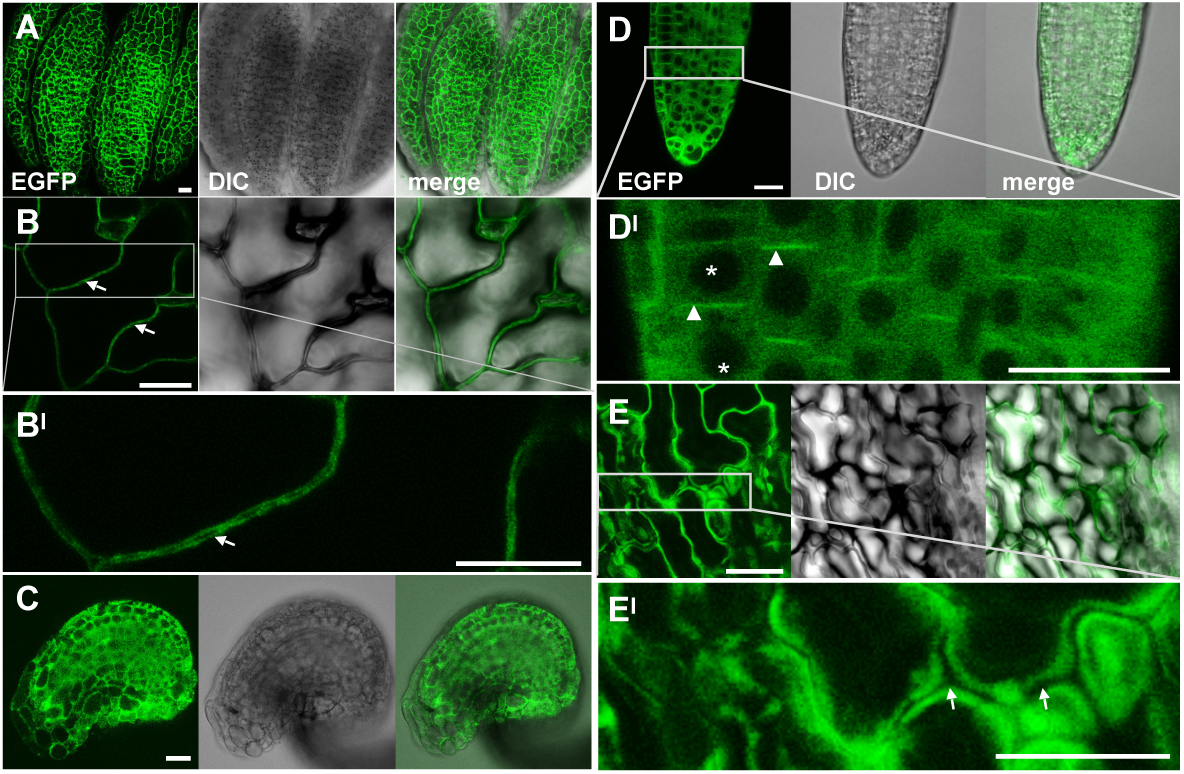
Subcellular localization of the pPDK1::PDK1.2:EGFP reporter signal in different tissues. Confocal micrographs are shown. (A) Mature anther. (B) Cauline leaf. (B^I^) higher magnification of the outlined region in (B). (C) Mature ovule. (D) Lateral root. (D^I^) higher magnification of the region in (D). (E) Sepal. (E^I^) higher magnification of the region in (E). (B, E^I^) Arrows indicate the absence of GFP signals at the cell wall, arrowheads indicate cortical or plasma membrane signal, and asterisks indicate absence of GFP signals in the nuclei. Scale bars: 20µm.

### PDK1 undergoes trans-autophosphorylation in vitro

To assess the in vitro phosphorylation activity of PDK1 we performed in vitro kinase assays using recombinant fusions of PDK1.1 and PDK1.2 to maltose binding protein (MBP:PDK1) or glutathione-S-transferase (GST:PDK1) that were produced in E. coli. In addition, we generated recombinant kinase-inactive versions of MBP:PDK1 (PDK1_KD_) by changing a conserved lysine to alanine in the catalytic domain of the two PDK1 homologs (MBP:PDK1.1_K73A_; MBP:PDK1.2_K74A_) (Fig 4A). We first investigated the in vitro autophosphorylation activities of MBP:PDK1.1 and MBP:PDK1.2. Confirming previous results we observed autophosphorylation of PDK1.1 and PDK1.2 (Fig 5B) [45–47, 51]. Next, we assessed if PDK1 undergoes trans-autophosphorylation. We observed that GST:PDK1.1 phosphorylated MBP:PDK1.2_KD_, and, vice-versa, GST:PDK1.2 phosphorylated MBP:PDK1.1_KD_ (Fig. 4B). The results indicate that PDK1.1 and PDK1.2 are active kinases that can physically interact and phosphorylate each other in vitro.

**Fig. 4.**
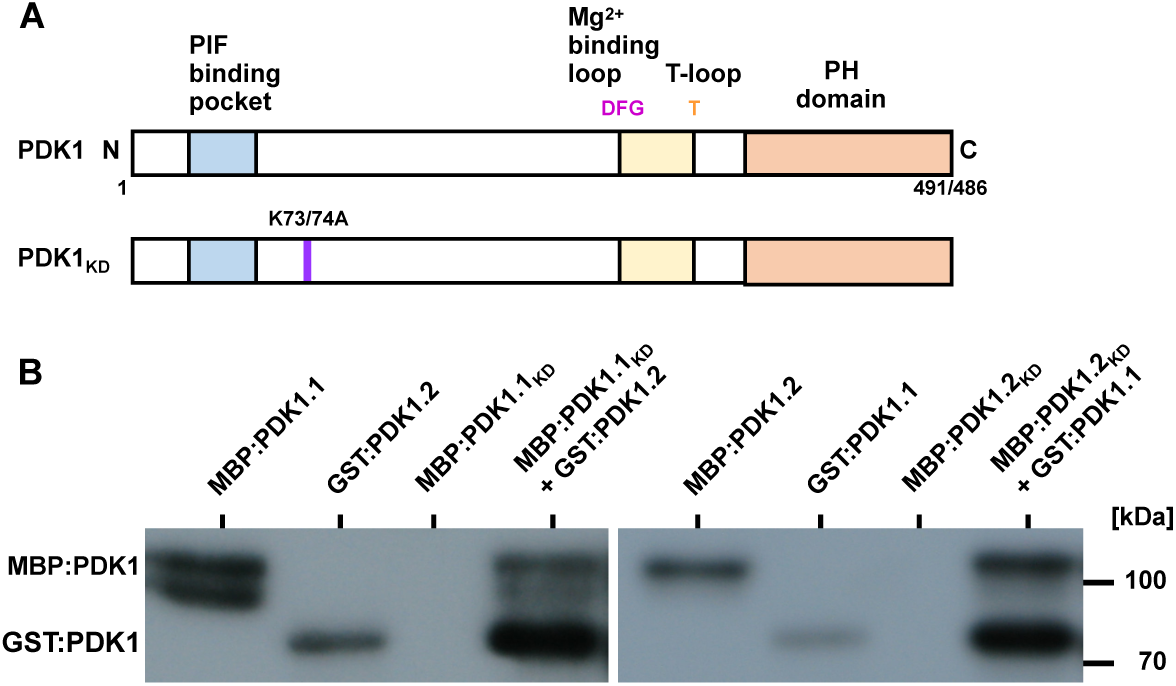
PDK1 in vitro kinase assays. (A) Cartoon outlining the domain structure of PDK1 and the various mutant versions. PDK1.1 has a length 491 and PDK1.2 of 486 amino acids, respectively. In PDK1_KD_, the conserved lysine residue at position 73 (in PDK1.1) or 74 (in PDK1.2) has been replaced by an alanine residue resulting in an inactive kinase. (B) Autoradiographs depicting in vitro kinase activity of MBP: and GST:PDK1 fusion proteins. Signal indicating auto-phosphorylation activity can be detected in all fusion proteins involving active MBP:PDK1. Note that MBP:PDK1_KD_ variants do not auto-phosphorylate but can be phosphorylated by active MBP:PDK1. Also note the trans-phosphorylation of MBP:PDK1.1KD or MBP:PDK1.2KD by GST:PDK1.2 or GST:PDK1.1, respectively.

**Fig. 5.**
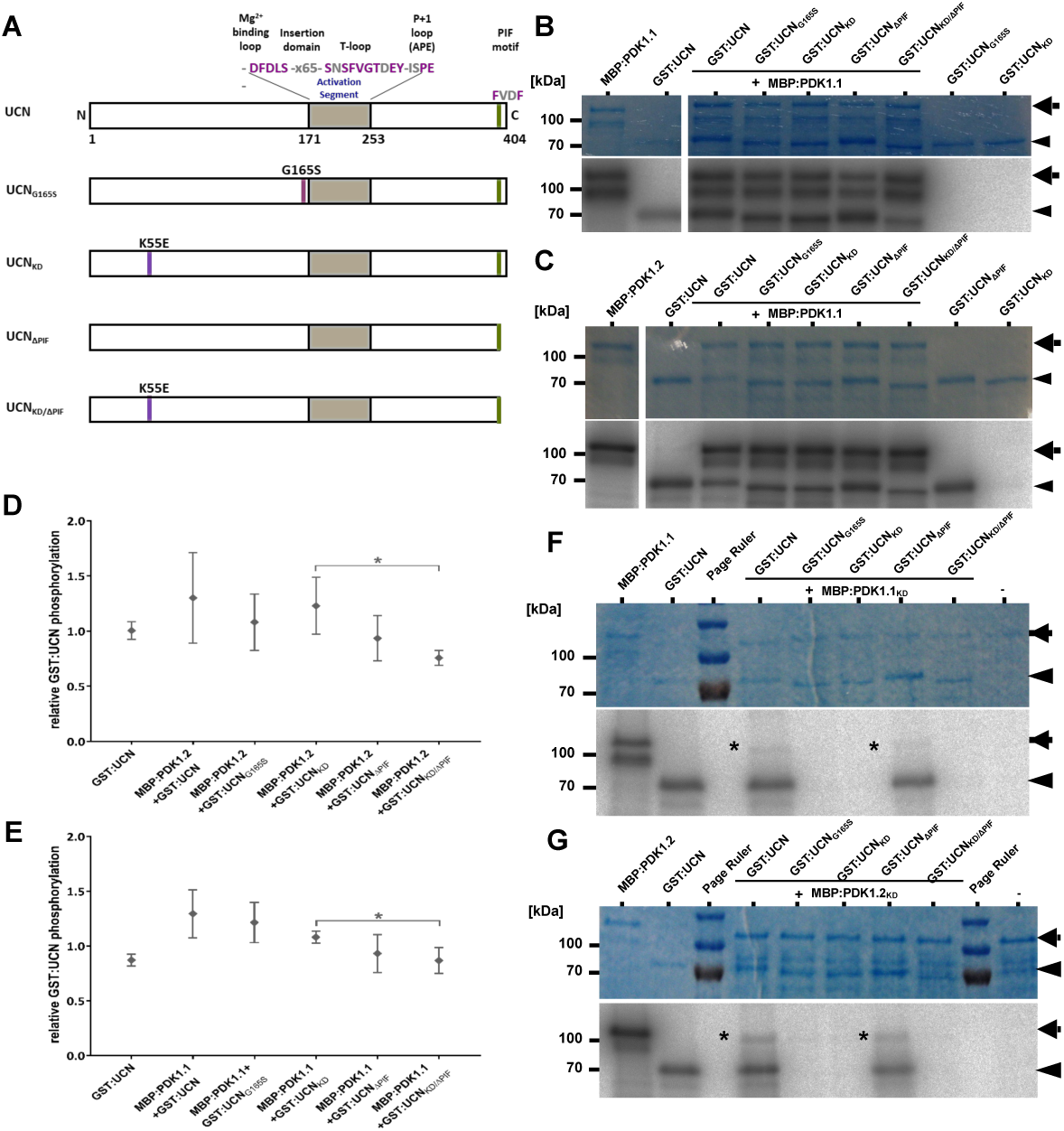
In vitro kinase assays with PDK1 in combination with different UCN variants. (A) Cartoon outlining the domain structure of UCN and marking the alterations in the different mutant variants. The plant AGC protein kinase specific insertion domain in the activation segment is also indicated. (B, C) and (F, G) Shown are coomassie brilliant blue (CBB)-stained gels (upper panel) and autoradiograms (lower panel) for MBP:PDK1.1, MBP:PDK1.2, MBP:PDK1.1_KD_, and MBP:PDK1.2_KD_, respectively, in combination with different GST:UCN versions. Arrows indicate the various MBP:PDK1 forms, arrowheads mark the different GST:UCN variants. (B) Kinase assays involving MBP:PDK1.1. Note the presence of corresponding labelled bands for all variants of GST:UCN. (C) Kinase assays involving MBP:PDK1.2. Note the presence of corresponding labelled bands for all variants of GST:UCN. (F) Kinase assays involving MBP:PDK1.1_KD_. Asterisk indicates MBP:PDK1.1_KD_ band weakly phosphorylated by GST:UCN or GST:UCN_ΔPIF_. (G) Kinase assays involving MBP:PDK1.2_KD_. Asterisk indicates MBP:PDK1.2_KD_ band weakly phosphorylated by GST:UCN or GST:UCN_ΔPIF_. (D, E) Intensity-based quantification of the bands indicated by the arrowheads and arrows in (B, C and F, G) (autoradiogram signal relative to the corresponding CBB gel band intensity) using ImageJ/Fiji [65, 66]. The results of three independent experiments (involving protein induction, purification and kinase assays) are shown. Please note statistically significant differences of phosphorylation intensity in a PIF motif-dependent manner (asterisks): (D) p = 0.031; (E) p = 0.039.

### PDK1 phosphorylates UCN in vitro

In a next step we tested if MBP:PDK1.1 and MBP:PDK1.2 can phosphorylate GST:UCN. To this end we performed in vitro kinase assays using wild-type and various mutant forms of recombinant MBP:PDK1 and GST:UCN fusion proteins. Four different types of mutant recombinant versions of UCN were generated (Fig 5A): UCNG165S (recapitulating the *ucn-1* defect, lacking kinase activity [9]), UCN_KD_ (UCN_K55E_, lacking kinase activity), UCN_ΔPIF_ (lacking the four C-terminal residues (FVDF) encompassing the predicted PIF domain), and UCN_KD/ΔPIF_ (lacking kinase activity and PIF domain).

We observed that MBP:PDK1.1 and MBP:PDK1.2 were able to trans-phosphorylate kinase-inactive GST:UCN variants (GST:UCNG165S, GST:UCN_KD_) (Fig 5B-D). Remarkably, and although there is a mild decrease in signal strength, we found that MBP:PDK1 can still phosphorylate GST:UCN variants lacking the PIF domain indicating that this fragment is not essential for PDK1/UCN interaction in vitro. We also performed the reciprocal experiment and tested if GST:UCN can phosphorylate MBP:PDK1 in vitro. Indeed, we observed that GST:UCN as well as GST:UCN_ΔPIF_ were able to phosphorylate MBP:PDK1.1_KD_ and MBP:PDK1.2_KD_ (Fig 5E-G). In both instances signal was relatively weak when compared to the signal obtained by GST:UCN autophosphorylation or phosphorylation of GST:UCN_KD_ by MBP:PDK1.1 or MBP:PDK1.2.

We further assessed if the presence of GST:UCN influenced the level of MBP:PDK1 activity. To this end titration experiments were performed in which constant levels of MBP:PDK1.1 or MBP:PDK1.2 were mixed with increasing amounts of GST:UCN or various mutant versions of GST:UCN (Fig 6). In both instances a slight but statistically significant decrease of MBP:PDK1 kinase activity was observed when increasing concentrations of GST:UCN were added to the reaction. A decrease in MPB:PDK1 kinase activity was not detected when kinase-defective variants of GST:UCN were used.

**Fig. 6.**
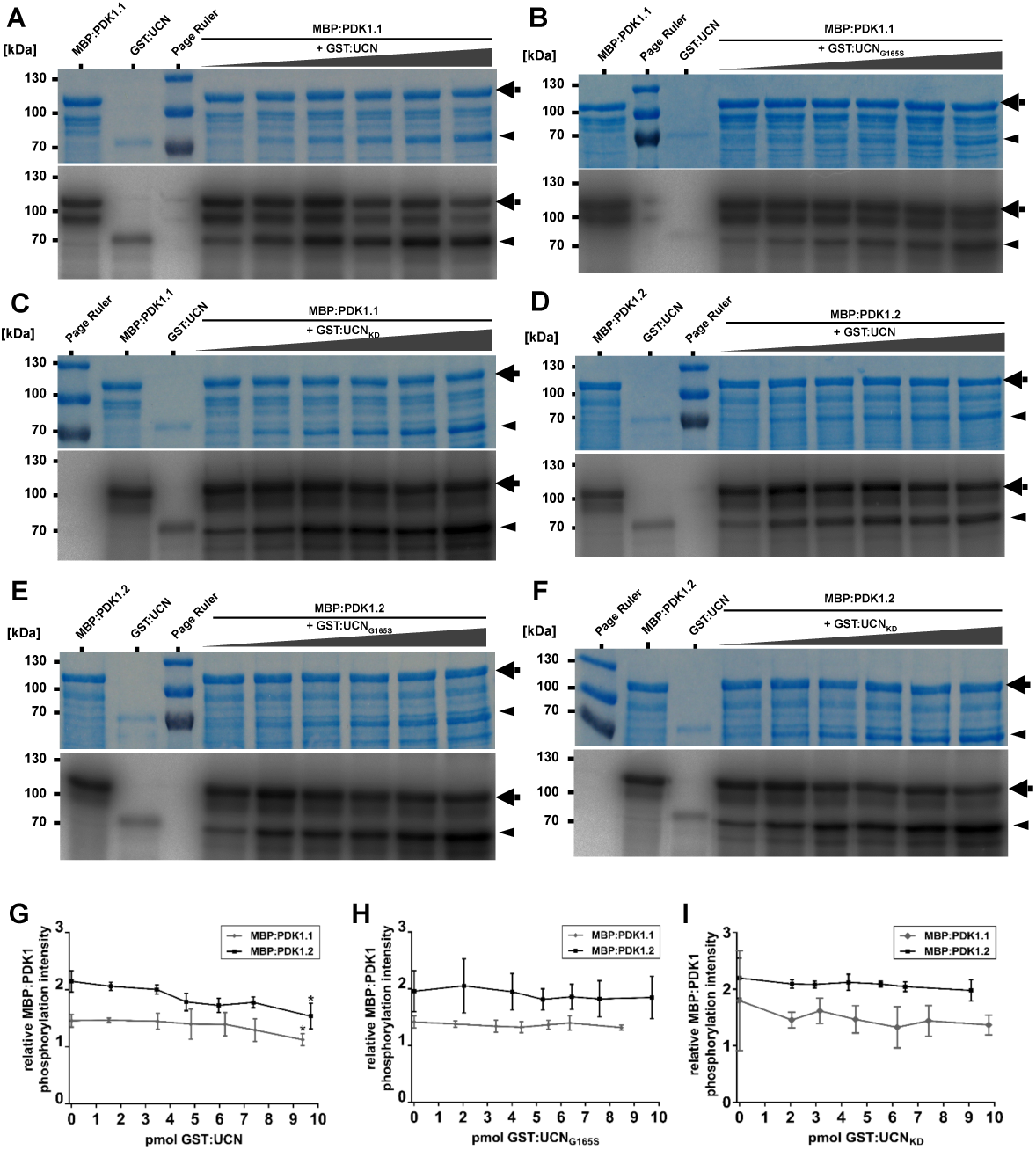
In vitro kinase assays with PDK1 and UCN. Coomassie brilliant blue (CBB)-stained gels and autoradiograms for MBP:PDK1.1 and MBP:PDK1.2, respectively, in combination with different GST:UCN versions are shown. Arrows indicate the MBP:PDK1 and arrowheads denote GST:UCN bands. (A-C) From left to right: First lane shows MBP:PDK1.1 and second lane shows GST:UCN autophosphorylation. Lanes 3 to 7 show combinations of constant levels of MBP:PDK1.1 with increasing amounts of GST:UCN, GST:UCNG165S and GST:UCN_KD_, respectively. (A) Note the slight reduction in MBP:PDK1 phosphorylation upon adding increasing amounts of GST:UCN. (D-F) Similar as in (A-C) but assays involve MBP:PDK1.2. (G-I) Intensity-based quantification of the bands indicated by the arrowheads and arrows in (A-F) (autoradiogram signal relative to the corresponding CBB gel band intensity) using ImageJ/Fiji. The results of three independent experiments (involving protein induction, purification and kinase assays) are shown. (G) Please note the significant decrease in relative MBP:PDK1 phosphorylation dependent on active UCN (asterisks). The p-values for the highest compared to the lowest level of UCN: MBP:PDK1: p = 0.021; MBP:PDK1.2: p = 0.024. (H-I) No decrease in relative MBP:PDK1 phosphorylation was observed in the presence of (H) UCN_KD_ or (I) UCN_G165S_. N = 3.

Taken together the results indicate that in in vitro assays PDK1 and UCN phosphorylate each other and that UCN can attenuate PDK1 activity.

### PDK1 and UCN interact in a plant cell

Next, we tested if PDK1 and UCN can interact in a living cell. We first undertook a yeast two-hybrid (Y2H) assay. We observed that UCN can interact with PDK1 in this system (Fig 7A). Moreover, we only detected interaction between wild-type proteins. We failed to observe interaction in Y2H assays involving variants carrying a mutation in the kinase domain of either UCN or PDK1, respectively, or exhibiting a deletion of PIF from UCN. Next, we investigated if PDK1 and UCN can interact in a living plant cell. Given the extremely low levels of *UCN* expression in the plant [9] we resorted to BiFC assays in Arabidopsis mesophyll protoplasts [52]. We observed BiFC signal in protoplasts co-transformed with *pSPYNE:PDK1.1* or *pSPYNE:PDK1.2* and *pSPYCE:UCN* constructs. (Fig 7B). Signal was apparently restricted to the cytoplasm correlating with the cytoplasmic localization of PDK1:EGFP in transgenic Arabidopsis lines. The result differs from similar BiFC experiments involving UCN and ATS where interaction was only seen in the nucleus [9]. Moreover, UCN can form homo-dimers in the cytoplasm and the nucleus [9]. As in the Y2H assay no BiFC-signal was observed when mutant UCN or PDK1 variants lacking either kinase activity or the PIF domain were employed (S4 Fig). The combined results indicate that UCN and PDK1 can interact in the cytoplasm of a plant cell, that kinase activity of UCN and PDK1 as well as the presence of the PIF domain in UCN are important for interaction in a living cell, and that UCN can interact with different partners in different subcellular compartments.

**Fig. 7.**
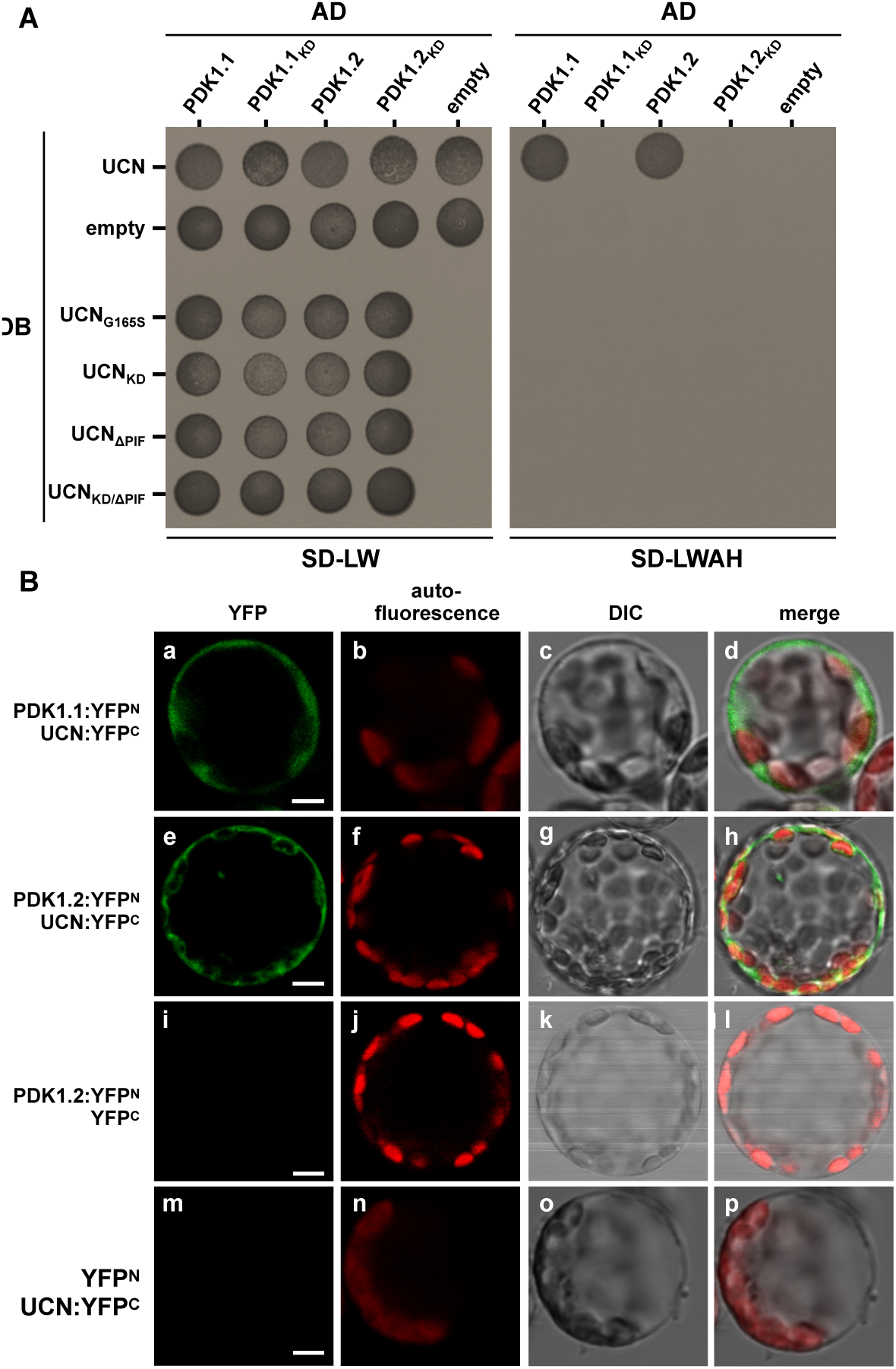
Yeast two-hybrid and BiFC assays. (A) Yeast two-hybrid assay. DB:UCN was tested with all four versions of PDK1, the other UCN versions were only tested with the wild type versions of PDK1. The left panel depicts yeast growth on transformation control plates (SD-LW), the right panel the interaction control plates (SD-LWH). Please note that only those yeast cells are able to grow on SDLWH, which were co-transformed with DB:UCN and AD:PDK1.1 or AD:PDK1.2. AD: activation domain of GAL4; DB: DNA-binding domain of GAL4; SD-LW: SD medium lacking Leu and Trp (transformation control); SD-LWAH: SD medium lacking Leu, Trp, Ade and His (interaction control). The experiment was repeated three times with similar results. (B) BiFC assay in Arabidopsis mesophyll protoplasts. The N-terminal part of YFP was fused to PDK1.1 or PDK1.2, respectively, and the C-terminal part of YFP was fused UCN. (a-d) Protoplasts co-transfected with PDK1.1:YFPN and UCN:YFPC, or (e-h) PDK1.2:YFPN and UCN:YFPC, respectively, show YFP fluorescence in the cytosol. 280 of 1566 protoplasts for PDK1.1 and 268 of 1543 for PDK1.2 showed YFP signals. (i-l) Protoplasts co-transfected with PDK1.2:YFPN and YFPC, or (m-p) YFPN and UCN:YFPC do not show any YFP signal (n > 1500). N = 3. Abbreviation: DIC, differential interference contrast. Scale bars: 5 µm.

### *UCN* is a negative regulator of *PDK1*

The results outlined above indicate that UCN and PDK1 physically interact in vitro and in plant cells. Next, we wanted to assess the biological relevance of such an interaction. To this end we performed a set of genetic analyses. We first investigated the phenotypes of *ucn-1 pdk1.1* and *ucn-1 pdk1.2* double mutants. Interestingly, we found in the *ucn pdk1* double mutants an essentially full restoration of the *ucn* ovule and flower phenotype to wild type (Fig 8, Table 1). The genetic result suggests that *UCN* is a negative regulator of *PDK1*. In *ucn* mutants elevated *PDK1* activity would lead to the mutant *ucn* phenotype which includes aberrant outgrowths on integuments and malformed petals [9]. In a double mutant the ectopic activity of *PDK1* would be absent resulting in a *pdk1*-like phenotype (which is apparently normal with respect to integument and petal development (Fig 8) [44].

**Table 1.** Characterization of *ucn*, *pdk1* and *ucn-1 pdk1* ovule phenotypes.

| Genotype | N protrusions |  |  |  | N<sup>a</sup> | % protrusions |
| --- | --- | --- | --- | --- | --- | --- |
|  | 0 | 1 | 2 | >2 |  |  |
| Col | 231 |  |  |  | 231 | 0 |
| *Ler* | 264 |  |  |  | 264 | 0 |
| *ucn-1* (*Ler*)<sup>b</sup> | 415 | 3303 | 546 | 89 | 4353 | 91 |
| *ucn-1* (Col)<sup>c</sup> | 1086 | 3384 | 316 | 39 | 4825 | 78 |
| *pdk1.1 ucn-1* | 8071 | 2009 | 0 | 0 | 10080 | 20 |
| *pdk1.2 ucn-1* | 9242 | 1133 | 0 | 0 | 10375 | 11 |
| *pdk1.1 pdk1.2* | 367 | 0 | 0 | 0 | 367 | 0 |
| *pdk1.1 pdk1.2 ucn-1* | 2130 | 241 | 0 | 0 | 2371 | 10 |

<sup>a</sup> Total number of ovules scored.
<sup>b</sup> original *ucn-1* allele in *Ler*.
<sup>c</sup> *ucn-1* introgressed into Col.

**Fig. 8.**
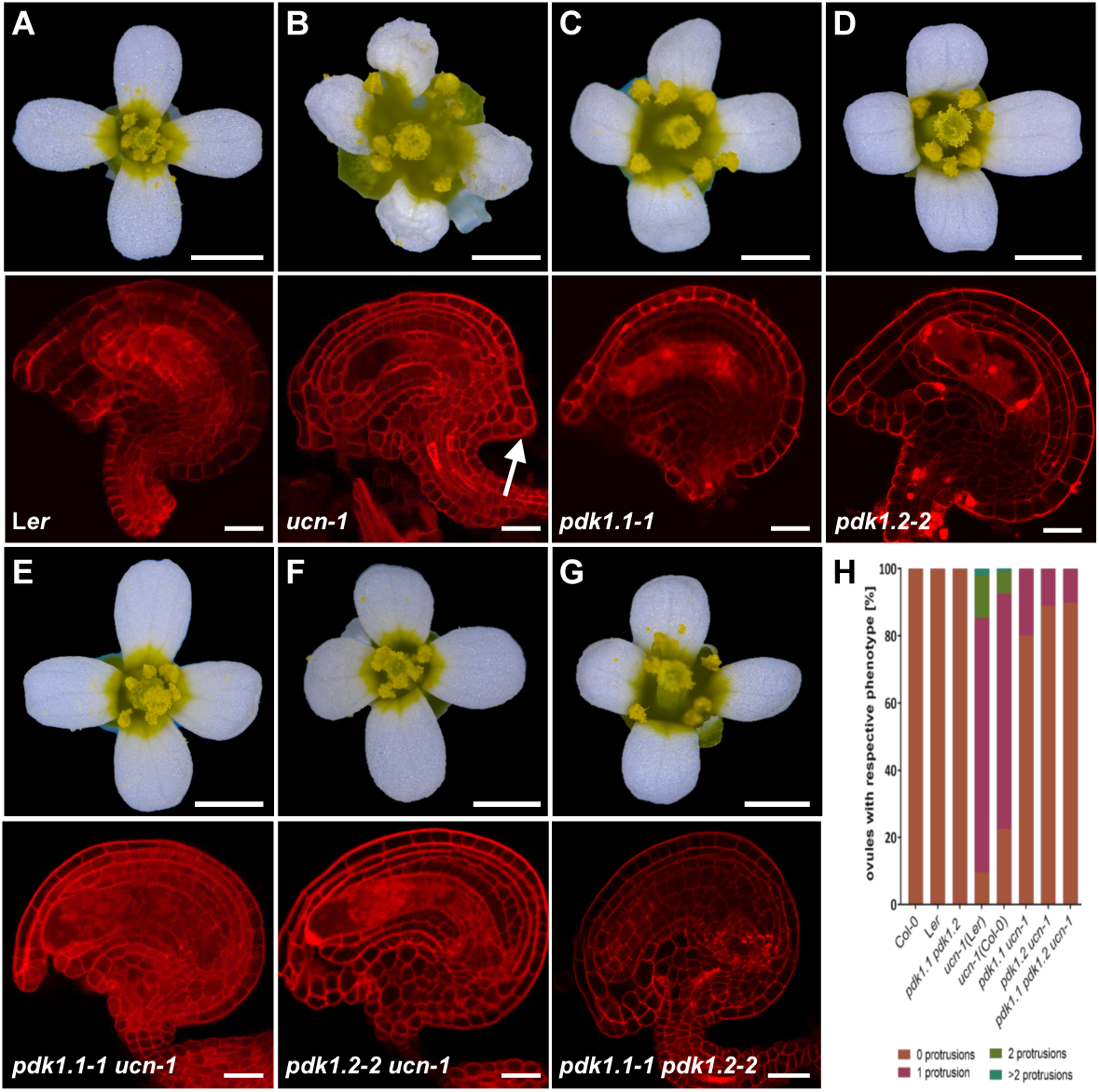
Analysis of *pdk1*, *ucn-1* and *pdk1 ucn-1* floral and ovule phenotypes. (A-F) Upper panel: Stage 13 flowers are shown (stages according to [67]). Lower panel: Confocal micrographs show about mid-optical sections through late stage 3 or stage 4 mPS-PI-stained ovules. Genotypes are indicated. (B) Upper panel: Note aberrant petal shape. Lower panel: arrow indicates ectopic protrusion. (C-G) Apparently normal phenotypes (compare with (A)). (H) Percentage of respective ovule phenotypes of L*er*, *ucn-1*, Col-0, *ucn-1* outcrossed to L*er* and Col-0 (F3 plants homozygous for *ucn-1*), respectively, and homozygous double mutants (*pdk1.1 ucn-1* and *pdk1.2 ucn-1*) and homozygous triple mutants (*pdk1.1 pdk1.2 ucn-1*). Knockouts of *pdk1.1*, *pdk1.2* or both restore the *ucn-1* phenotype to 73 % and 90 % of the WT level, respectively (*ucn-1 pdk1.1-2*: 73%; *ucn-1 pdk1.1-1*: 80%; *ucn-1 pdk1.2-3*: 86%; *ucn-1 pdk1.2-2*: 90%). Sample sizes are given in Table 1. Scale bars: (A-F) Upper panels, 1mm; lower panels, 20 µm.

If the notion of *UCN* being a negative regulator of *PDK1* function was valid one would expect ectopic activity of *PDK1* to result in a *ucn*-like phenotype. To test this assumption, we generated EGFP fusions of PDK1.1 and PDK1.2 under the control of the UBIQUITIN promoter (pUBQ10, At4g05320) and transformed wild-type L*er* plants with the corresponding transgenes. In both cases we investigated the phenotypes of 11 independent transgenic T2 lines homozygous for the transgene. In nine pUBQ::PDK1.1:EGFP lines we observed distorted petals and ovules with protrusions indicating that overexpression of *PDK1* results in a *ucn*-like phenocopy (Fig 9, Table 2). Similar phenotypes were observed in eight pUBQ::PDK1.2:EGFP lines. We also crossed two of the phenotypic pUBQ::PDK1.1:EGFP and pUBQ::PDK1.2:EGFP lines into *ucn-1*. In all instances we observed an increase in the number and size of integumentary protrusions in *ucn-1 pUBQ::PDK1:EGFP* when compared to protrusions formed in *ucn-1*. These data indicate that ectopic expression of *PDK1* in a *ucn-1* background aggravates the *ucn-1* phenotype further. The data support the notion of *UCN* being a negative regulator of *PDK1* and hint at the presence of additional, as yet unidentified, repressors of *PDK1*.

**Table 2.**
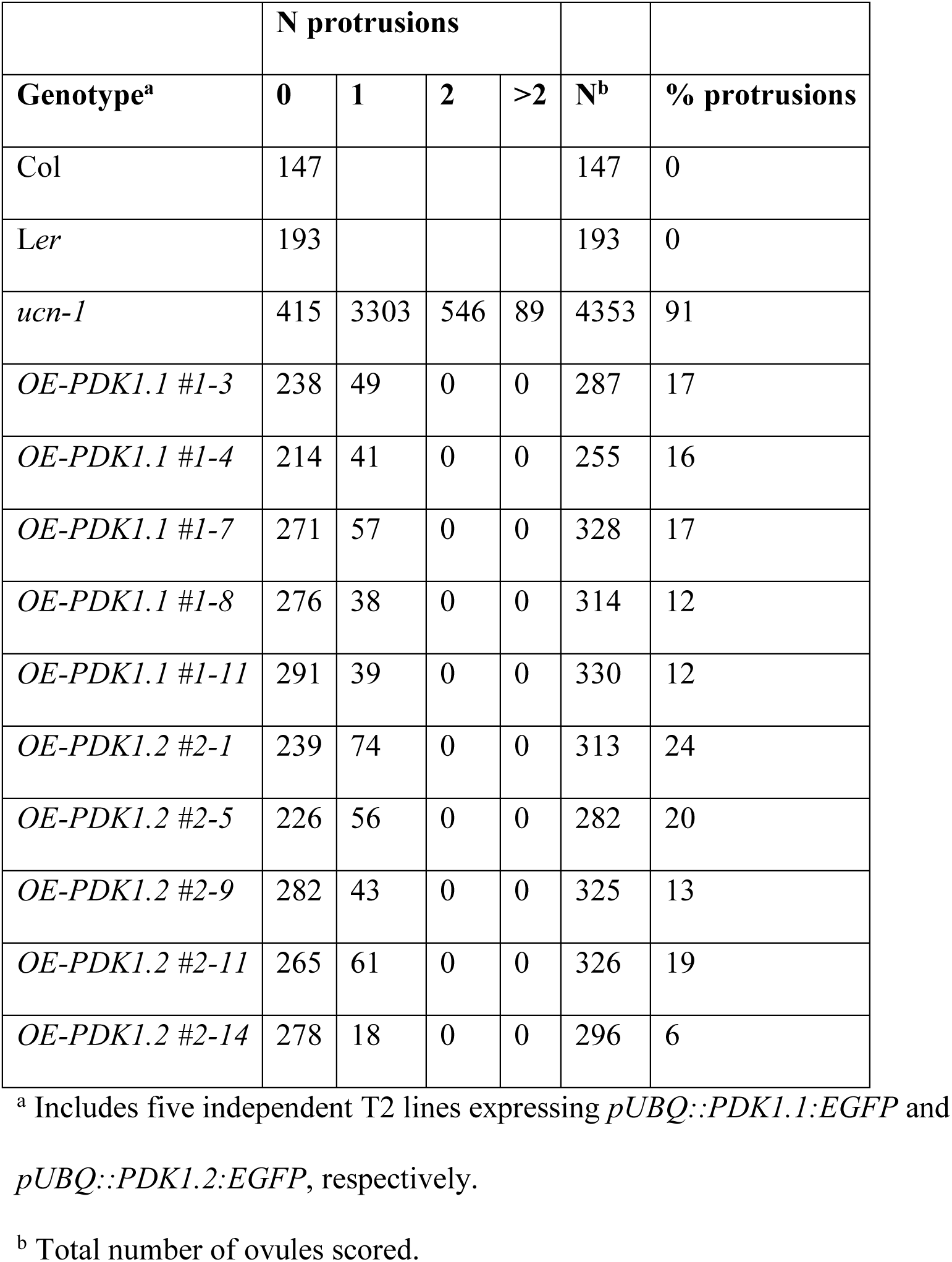
Ovule phenotypes of *PDK1* overexpressing lines.

**Fig. 9.**
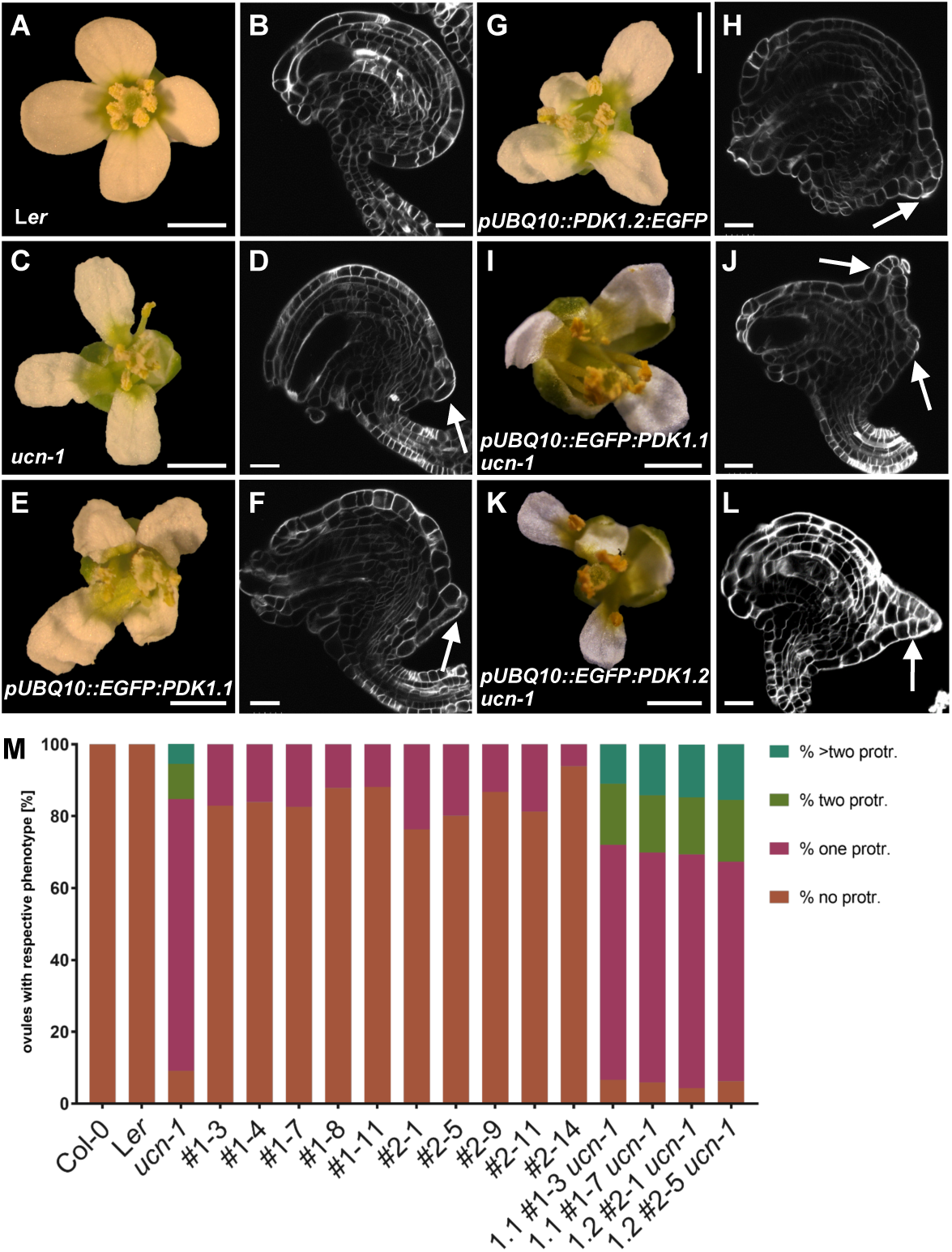
Phenotypes of *pUBQ::PDK1:EGFP* lines. (A, C, E, G, I, K) Stage 13 flowers are shown. (B, D, F, H, J, L) Confocal micrographs show about mid-optical sections through late stage 3 or stage 4 SCRI Renaissance 2200-stained ovules. Genotypes are indicated. (M) Percentage of respective ovule phenotypes in WT, *ucn-1*, and ten independent T-DNA lines overexpressing PDK1:EGFP (#2-1 to #2-14 overexpressing *PDK1.2:EGFP*, and #1-3 to #1-11 overexpressing *PDK1.1:EGFP*). Sample sizes are given in Table 2. (D, F, J, L) Arrows indicate protrusions. Scale bars: (A, C, E, G, I, K) 1mm; (B, D, F, H, J, L) 20 µm.

The finding that ectopic *PDK1* expression results in a *ucn*-like phenotype raises the possibility that *UCN* could function as a transcriptional regulator of *PDK1*. To test this hypothesis, we assessed *PDK1.1* and *PDK1.2* expression in *ucn-1* ovules by whole-mount ISH. No obvious expression differences could be detected between wild type and *ucn-1* (Fig 10A-D). We also assessed *PDK1.1* and *PDK1.2* transcript levels in wild type and *ucn-1* flowers and stems by quantitative real-time PCR (qPCR) (Fig 10E). We could not observe a significant difference in *PDK1.1* or *PDK1.2* transcript levels between wild type and *ucn-1*. These observations indicate that the negative regulation of *PDK1* by *UCN* occurs at the post-transcriptional level.

**Fig. 10.**
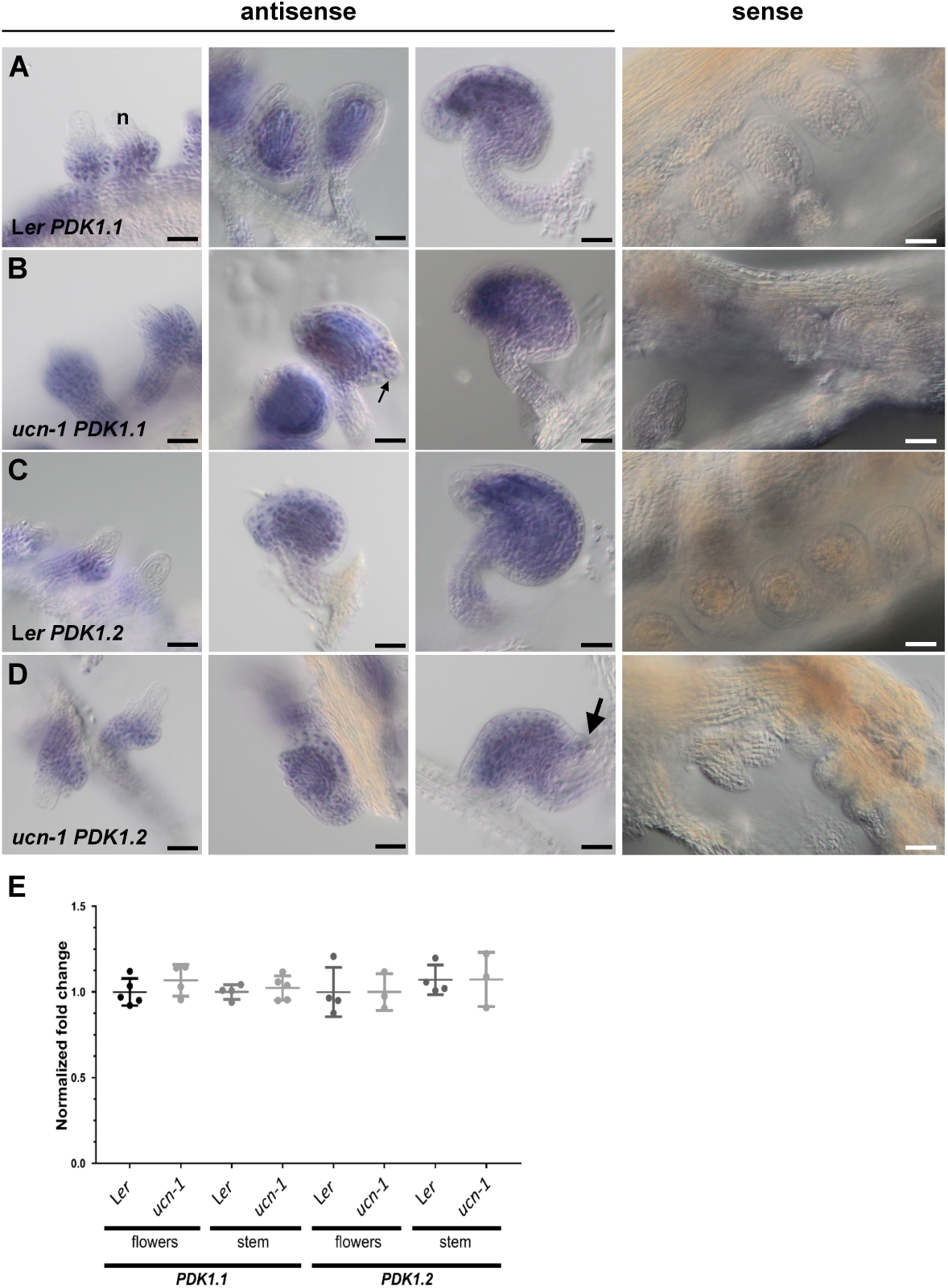
*PDK1* expression in *ucn-1*. (A-D) Whole-mount in situ hybridization. Genotypes and probes are indicated. Left three panels: stage 2-III/IV ovules (left), stage 3-IV/V (center) and stage 3-VI/4-I ovules. There is no apparent difference in the expression pattern between the genotypes. The sense control involves ovules of different stages. No signal above background is detected. (E) qPCR analysis using RNA isolated from flower (stages 1-13) and stems. Genotypes and probes are indicated. Between three to five biological replicates were used for each tissue and genotype. Means ± SD are shown. Expression levels for *PDK1.1* or *PDK1.2* do not noticeably vary between genotypes. At2g28390, At4g33380, and At5g46630 were used as reference genes [9]. p-values (L*er* vs. ucn-1) are as follows: *PDK1.1* flowers, p = 0.28; *PDK1.1* stem, p = 0.56; *PDK1.2* flowers, p = 0.44; *PDK1.2* stem, p = 0.55. Scale bars: 25 µm.

### *PDK1* is required for *ucn*-like aberrant integumentary growth caused by overexpression of *ATS*

The model put forward for the functional relationship between PDK1 and UCN has strong similarities to a scenario that was proposed for the UCN/ATS interaction [9]. UCN/ATS protein interaction appears to be restricted to the suppression of ectopic outgrowths in integuments as other aspects of the *ucn* phenotype, such as malformed petals, were unaffected in *ucn ats* double mutants [9]. Moreover, BiFC experiments indicate that complex formation between UCN and ATS occurs in the nucleus not the cytoplasm. Thus, the available data suggest that the UCN attenuation of PDK1 is of broader importance than the inhibition of ATS activity by UCN.

These considerations raise the question how PDK1 and ATS relate to each other. We first tested if PDK1 can phosphorylate ATS in vitro. We could detect phosphorylation of a recombinant translational fusion of thioredoxin to ATS by MBP:PDK1 in in vitro kinase assays (S5A Fig). However, signal intensity was very low in comparison to for example phosphorylation of GST:UCN_KD_ by MBP:PDK1. We next assessed if PDK1 can interact with ATS in a plant cell. We did not observe PDK1/ATS interaction in BiFC experiments indicating that the PDK1/ATS interaction observed in vitro does not occur in a plant cell (S5B Fig). In line with this view, our data suggest that PDK1 is present in the cytoplasm but excluded from the nucleus. By contrast, we observed interaction between UCN and ATS in BiFC assays in the nucleus as expected for a transcription factor [9]. To further assess the role of PDK1 in ATS function we asked whether *PDK1* is required for *ucn*-like ectopic outgrowth formation in *sk21-D* plants. In the activation tagging mutant *sk21-D* [53] *ATS* transcript levels are elevated about 45 fold compared to wild type but its spatial expression remains normal [9]. Interestingly, *pdk1.1 sk21-D* or *pdk1.2 sk21-D* double mutants failed to produce *ucn-*like integumentary protrusions (Fig 11A-D). In addition, we performed the complementary experiment and tested if *ATS* is required for the formation of integumentary protrusions in *pUBQ::PDK1:EGFP* lines. To this end we crossed two independent pUBQ::PDK1.1:EGFP and pUBQ::PDK1.2:EGFP lines into *ats-3*. We observed that ovules of *ats-3 pUBQ::PDK1.1:EGFP* or *ats-3 pUBQ::PDK1.2:EGFP* plants did not generate ectopic outgrowths (Fig 11E-G). The observations indicate that *PDK1* activity is required for integumentary protrusion formation occurring in plants with abnormally elevated *ATS* expression. Furthermore, activity of only one of the two *PDK1* loci is insufficient to provide enough *PDK1* activity to permit aberrant growth formation caused by too high levels of *ATS* activity. At the same time, *ATS* is necessary for integumentary protrusion formation in lines overexpressing *PDK1:EGFP*.

**Fig. 11.**
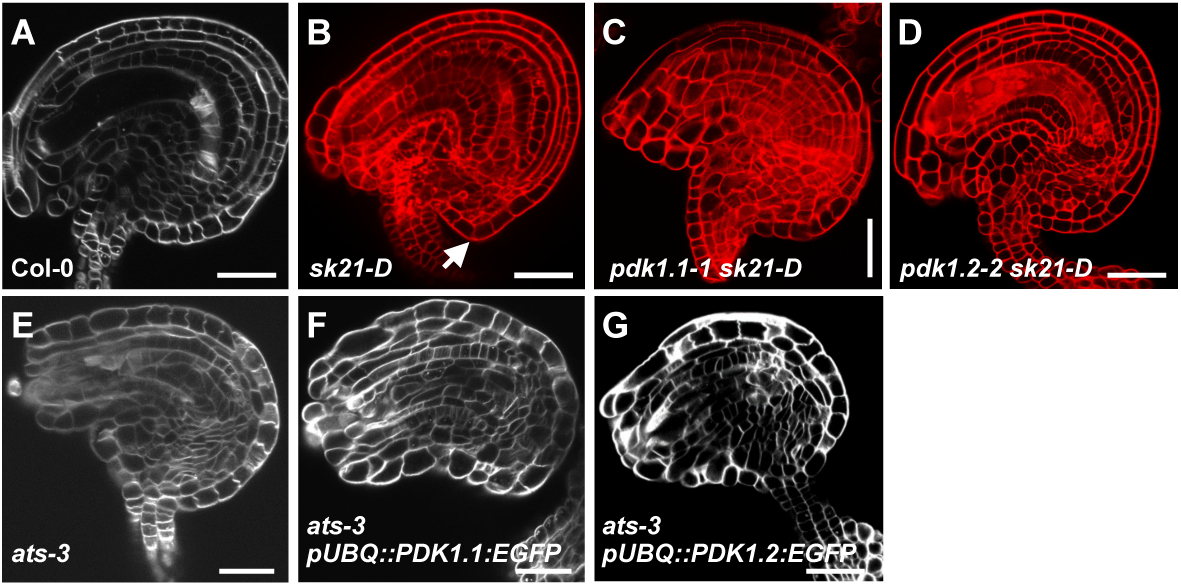
Analysis of the interaction between *PDK1* and *ATS*. Confocal micrographs show about mid-optical sections through late stage 3 or stage 4 ovules. (A, E-G) SCRI Renaissance 2200-stained ovules. (B-D) mPS-PI-stained ovules. Genotypes are indicated. (B) Arrow indicates protrusion. (C, D) and (F, G) Note absence of protrusions. Sample sizes are given in Table 3. Scale bars: 20 µm.

**Table 3.** Characterization of *ats*-related ovule phenotypes.

|  | **N protrusions** |  |  |  |
| --- | --- | --- | --- | --- |
| **Genotype** | **0** | **1** | **N<sup>a</sup>** | **% protrusions** |
| *sk21D* | 188 | 151 | 339 | 45 |
| *pdk1.1 sk21D* | 359 | 8 | 376 | 2 |
| *pdk1.2 sk21D* | 379 | 5 | 384 | 1 |
| *ats-3* | 167 | 0 | 167 | 0 |
| *ats-3 OE-PDK1.1 #3* | 309 | 4 | 313 | 1 |
| *ats-3 OE-PDK1.2 #1* | 317 | 3 | 320 | 1 |

<sup>a</sup> Total number of ovules scored.

## Discussion

Plant *PDK1* has been implied in several stress responses, including the control of cell death, the response to fungal pathogenic elicitors, or basal disease resistance [37, 38, 40, 41]. Here, we focused on the potential involvement of *PDK1* in growth control and signaling mediated by the AGCVIII protein kinase UCN.

In the current model of the regulation of AGC protein kinases the master regulator PDK1 activates a set of different downstream AGC protein kinases and plays a central role in growth control [23, 36]. In plants, the situation appears to be more complex. While in tomato down-regulation of *PDK1* results in lethality [38] complete absence of *PDK1.1* and *PDK1.2* function only leads to minor growth defects in Arabidopsis [44] (this study). Thus, reduced or absent levels of *PDK1* function can be relatively easily accommodated at least when grown under optimal conditions. It indicates that other kinases must have assumed essential growth functions that are carried out by animal PDK1, at least in Arabidopsis and possibly in other plant species as well [40, 43].

Our analysis using several functional PDK1:EGFP reporters indicates that Arabidopsis PDK1 localizes to the cytoplasm. In cells of the root meristem and transition zone, however, PDK1 appears to be present at the PM as well. This subcellular distribution resembles data from animal cells which indicate that PDK1 is found predominantly in the cytoplasm of unstimulated cells. A noticeable fraction of PDK1 at the PM can only be detected after application of for example growth factors [29, 32, 33].

The results obtained from in vitro phosphorylation studies as well as Y2H and BiFC assays are compatible with the notion that PDK1 and UCN physically interact in vitro and in plant cells. Interestingly, we observed in vitro interactions between PDK1 and UCN, as monitored by kinase assays, even in the presence of deletions of the PIF domain of UCN or the absence of kinase activity of either PDK1.1, PDK1.2, or UCN. By contrast, we could only detect interactions in living cells, as assessed in Y2H and BiFC assays, when we employed wild-type UCN or PDK1 but not the various tested mutant versions. A lack of accordance between different types of assays was also observed when for example investigating the interactions between PDK1 and OXI1/AGC2-1, an AGCVIII protein kinase [46] involved in oxidative burst responses and growth promotion [39, 44, 54]). These results reinforce the notion that interaction data obtained from in vitro experiments may not always correlate with results based on assays involving living plant cells.

It is not entirely clear what is the ultimate biochemical cause for our failure to observe interaction between PDK1 and UCN in Y2H or BiFC assays employing the tested mutant versions of both PDK1 homologs and UCN. Alterations of the migration pattern in denaturing protein gels of mutant versus wild-type versions of UCN or PDK1 fusion proteins may hint at conformational changes that impairs interaction. In the end, however, our findings that the tested mutant versions of UCN and PDK1 do not interact in BiFC assays in plant cells appear to be functionally relevant as they are in line with the observed genetic interaction between *PDK1* and *UCN*.

Our genetic data support a functional relevance of the interactions between PDK1 and UCN detected in vitro, in yeast, and in protoplasts. However, it is unlikely that the observed in vitro phosphorylation of GST:UCN by MPB:PDK1 reflects an essential activation of UCN by PDK1 in planta. If this notion was true one would expect a *ucn-*like phenotype in *pdk1.1. pdk1.2* double mutants. We did not observe such a phenotype, however, we cannot exclude that another, as yet unidentified, protein kinase substitutes for PDK1. In any case, the observed rescue of the *ucn* integumentary outgrowths and petal malformations in *pdk1.1 ucn-1* or *pdk1.2 ucn-1* double mutants suggests *UCN* to act as a negative regulator of *PDK1*. As we did not detect altered *PDK1* transcript levels in *ucn* mutants we propose that the regulation occurs at the post-transcriptional level. This notion is also supported by the observation that increasing concentrations of GST:UCN mediated a mild decrease in MBP:PDK1 in vitro kinase activity. Interestingly, eliminating the function of only one of the two *PDK1* homologs was sufficient to achieve suppression of the *ucn* phenotype. It indicates that total level of combined *PDK1.1* and *PDK1.2* activity and/or stoichiometry in a PDK1-containing protein complex may be important aspects of the *UCN*-*PDK1* interaction. A dynamic equilibrium between monomeric and dimeric forms is believed to be important for the function of PDK1 in animal cells [32, 33].

We observed *ucn*-like integumentary outgrowths and petal malformation in *pUBQ::PDK1:EGFP* lines indicating that overexpression of *PDK1* interferes with growth regulation during development of these tissues. Thus, it seems that ectopic activity of *PDK1* must be avoided to allow proper tissue morphogenesis. In a parsimonious interpretation of our results we postulate that UCN inhibits PDK1 activity through direct protein-protein interactions in the cytoplasm. In this scenario UCN attenuates PDK1 activity to keep PDK1 activity below a certain threshold. Given the extremely low expression levels of *UCN* found in ovules and flowers [9] we suggest that overexpression of *PDK1* results in high PDK1 protein levels that titrate out available UCN proteins. Elevating PDK1 activity beyond the threshold would therefore lead to a deregulation of growth control in integuments and petals. Interestingly, overexpression of *PDK1* is an important feature of many human tumors [26]. We propose that UCN is part of the mechanism that attenuates PDK1 function in Arabidopsis and thus prevents the deregulation of growth control in integuments and petals.

Previous results [9, 14] and the data presented here are compatible with the notion that UCN attenuates the activity of at least two proteins with diverse functions, the protein kinase PDK1 and the putative transcription factor ATS. The effects of the interaction between UCN and ATS appear to be restricted to the regulation of planar growth in integuments [9, 14]. By contrast, the results shown here indicate that the interaction between UCN and PDK1 is of broader relevance and controls integument and petal development. How does PDK1 relate to ATS? It is unlikely that PDK1 directly affects ATS. We observed only very weak phosphorylation of ATS by PDK1 in in vitro kinase assays. Moreover, BiFC experiments did not support physical interaction between PDK1 and ATS in a plant cell and our results failed to provide evidence for a presence of PDK1 in the nucleus. In addition, *PDK1* does not seem to be involved in the promotion or inhibition of *ATS* activity as *pdk1.1. pdk1.2* double mutants did not show an *ats* or *sk21-D*-like phenotype. Thus, the in vitro phosphorylation data may not be relevant in vivo. However, ectopic outgrowth formation in integuments upon overexpression of *PDK1* or *ATS* depended on the presence of *ATS* and *PDK1*, respectively. Thus, the present data support the view that PDK1 may represent a more globally acting factor that conditions a cellular context in which for example exceedingly high levels of ectopic ATS activity can exert its detrimental effects on growth regulation in integuments. In this model UCN functions at the nexus of two separate pathways and balances the activity of more global as well as local effectors of growth control. UCN thereby maintains the correct growth patterns underlying integument and petal morphogenesis. So far, the prevailing model of the regulation of AGC protein kinases states that the master regulator PDK1 activates a set of different downstream AGC protein kinases [23, 36]. The finding that an AGC protein kinase attenuates PDK1 is a new finding not just for plants but also for other eukaryotes. Thus, apart from enhancing our understanding on the control of planar growth in plants, this work expands the general conceptual framework of AGC protein kinase regulation in eukaryotes.

## Materials and Methods

### Plant work, plant genetics and plant transformation

Arabidopsis thaliana (L.) Heynh. var. Columbia (Col-0) and var. Landsberg (erecta mutant) (Ler) were used as wild-type strains. Plants were grown as described earlier [55]. The *ucn-1* mutant (in L*er*) was described previously [9], and *pdk-1* T-DNA lines were described before [44]. T-DNA insertion lines were received from the NASC (pdk1.1-1 SALK_053385, pdk1.1-2 SALK_113251, pdk1.2-2 SAIL_62_G04, pdk1.2-3 SAIL_450_B01). Wild-type and *pdk1.1 pdk1.2* mutant plants were transformed with different constructs using Agrobacterium strain GV3101/pMP90 [56] and the floral dip method [57]. Transgenic T1 plants were selected on Hygromycin (25 µg/ml) plates and transferred to soil for further inspection. Gene identifiers: *ATS* (At5g42630), *PDK1.1* (At5g04510), *PDK1.2* (At3g10540), *UCN* (At1g51170).

### Recombinant DNA work

For DNA and RNA work standard molecular biology techniques were used. PCR-fragments used for cloning were obtained using Phusion or Q5 high-fidelity DNA polymerase (both New England Biolabs, Frankfurt, Germany). All PCR-based constructs were sequenced. The Gateway-based (Invitrogen) pDONR207 was used as entry vector, and destination vectors pMDC43 and pMDC83 [58] were used as binary vectors. Detailed information for all oligonucleotides used in this study is given in S1 Table. The kinase-deficient mutant versions of either PDK1 or UCN were generated by site-directed mutagenesis approaches. The conserved lysine residues at positions 73 (PDK1.1), 74 (PDK1.2) or 55 (UCN) were replaced with alanine residues (PDK1) or a glutamic acid residue (UCN), respectively. UCNG165S was generated in a similar approach by replacing the Gly165 residue by a serine.

### PCR-based gene expression analysis

Floral tissue for quantitative real-time PCR (qPCR) was harvested from plants grown under long day conditions. With minor changes, tissue collection, RNA extraction and quality control were performed as described previously [59]. cDNA synthesis, qPCR, and analysis was done essentially as described [9].

### Reporter constructs

For plasmid pPDK1.2::gPDK1.2:EGFP pMDC83, 4.253 kb of gPDK1.2 sequence was amplified from Col-0 genomic DNA including promoter sequence spanning genomic DNA up to the 3′ end of the adjacent gene (1.245 kb) and 3′UTR of PDK1.2 and cloned into pDONR207. pMDC83 (35S promoter was removed) was used as destination vector. For overexpression constructs, gPDK1.1 or gPDK1.2 were amplified from Col-0 genomic DNA and cloned into pDONR207. As destination vectors, pMDC43 or pMDC83 (35S promoter was replaced by either pUBQ10 or p16) were used, respectively.

### Generation, expression and purification of recombinant proteins

UCN and PDK1 coding sequences were amplified from floral cDNA (Ler) and cloned into pGEX-6P-1 (GE Healthcare, Munich, Germany) or pMal-c2x (New England Biolabs, Frankfurt, Germany). The clones were expressed in the E. coli strain BL21 (DE3) pLysS. Expression from the pGEX vector leads to proteins fused to a N-terminal Glutathione Transferase (GST) protein, expression from pMal leads to proteins fused to a N-terminal Maltose Binding Protein (MBP). The ATS coding sequence was cloned into pET32a (Novagen) leading to an N-terminal 6xHis:Thioredoxin (Trx) fusion protein. For protein expression and purification, bacterial cultures were grown to OD 0.6-0.8 at 37°C. Then, the bacteria were induced with 0.8 mM isopropyl-beta-thio galactopyranoside (IPTG) for UCN, 1.5 mM IPTG for PDK1, and 0.5 mM IPTG for ATS, and grown at 30°C for 4 h. Subsequently, the recombinant proteins were purified from the bacteria by batch purification under native conditions using the Glutathione Sepharose 4 Fast Flow (GE Healthcare) for GST fusion proteins, amylose resin (New England Biolabs) for MBP fusion proteins, and Protino® Ni-TED packed columns for 6xHis:Trx fusion proteins according to the manufacturer’s instructions. For PDK1, four different protein versions were expressed and purified (PDK1.1 WT, PDK1.1_KD_ (kinase deficient version containing a K73A mutation), PDK1.2 WT and PDK1.2_KD_ (kinase deficient version containing a K74A mutation)). For UCN, five different protein versions were expressed and purified (UCN WT, UCNG165S (*ucn-1* mutation, kinase deficient), UCN_KD_ (kinase deficient version containing a K55E mutation), UCN_ΔPIF_ (kinase active version lacking the PIF motif at the C-terminus), and UCN_KD/ΔPIF_ (kinase deficient version lacking the PIF motif at the C-terminus)).

### In vitro kinase assays

For kinase assays, the proteins were purified as described above and concentrations were estimated on a 12% SDS PAGE using BSA as standard protein. Assays were performed with approximately 500 ng of respective protein(s) and incubated in HMK buffer (10 mM HEPES, 10 mM MgCl_2_, 10 µM ATP and 2 µCi either γ-^32^P-ATP or γ-^33^P-ATP (Hartmann Analytik, Braunschweig, Germany)) at RT for 1 h. Reactions were stopped by adding 4 µL 6xLaemmli buffer and boiling at 95°C for 5 min. In order to separate the proteins, 12% SDS PAGE was performed. Subsequently, the gels were stained with Coomassie Brilliant Blue G250, destained in 10% acetic acid and dried. Phosphorimager plates were exposed at RT over night and signals were detected using a Fuji BAS Phosphorimager (Fujifilm, Düsseldorf, Germany).

### Yeast two-hybrid assays

For yeast two-hybrid assays, the above-mentioned four PDK1 and five UCN versions were used. The coding sequences of these versions were cloned into pGBKT7 and pGADT7 vectors, respectively (Clontech Laboratories/Takara Bio, Saint-Germain-en-Laye, France). Plasmids were transformed into yeast strain AH109 and transformants were selected on SD_-LW_ medium (SD medium without Leu and Trp). Three independent colonies of each combination were resuspended in 500 µL ddH_2_O, diluted 1:100 and 10µL of the dilutions were spotted on SD_-LWHA_ (SD medium without Leu, Trp, His and Adenine) supplemented with 5 mM 3-AT and grown at 30°C for 3 days.

### BiFC assays

For Bimolecular Fluorescence Complementation (BiFC), the above-mentioned four PDK1 and five UCN versions were cloned into pSPYCE-35S and pSPYNE-35S vectors, respectively [52]. Plasmids were transformed into Col-0 mesophyll protoplasts as described [60]. Protoplasts were incubated gently shaking at 21°C in the dark for 10 to 15 hours and subsequently imaged using a FV1000 confocal microscope (Olympus, Hamburg, Germany) with excitation at 515 nm and detection at 521 – 559 nm.

### In situ hybridization and microscopy

Whole mount in situ hybridization of ovules was performed essentially as described [61]. Digoxigenin-labeled probe generation has been described earlier [62]. An Olympus BX61 upright microscope with DIC optics was used for microscopic analysis of the slides. Confocal laser scanning microscopy (CSLM) using modified pseudo-Schiff propidium iodide (mPS-PI) staining or detection of EGFP was performed as described earlier [63]. The SCRI Renaissance 2200 staining protocol was carried out as described [64].

## Acknowledgments

We acknowledge Michael Papacek and Moritz Ruschhaupt for technical help. We also thank Ajeet Chaudhary, Ulrich Z. Hammes, and Prasad Vaddepalli for critical reading of the manuscript. This work was funded by the German Research Council (DFG) through grant SCHN 723/7-1 to KS.

## Supporting information

**S1 Fig. Protein alignment of PDK1.1, PDK1.2, and At2g20050.**

**S2 Fig. Restoration of *pdk1.1 pdk1.2* phenotype by transgenes expressing translational fusions of PDK1 to EGFP.**

**S3 Fig. Subcellular localization of pUBQ::PDK1:EGFP reporter signals.**

**S4 Fig. BiFC assays of PDK1 with mutant versions of UCN.**

**S5 Fig. PDK1 phosphorylates ATS in vitro.**

**S1 Table. Primers used in this study.**

**S1 Fig.**
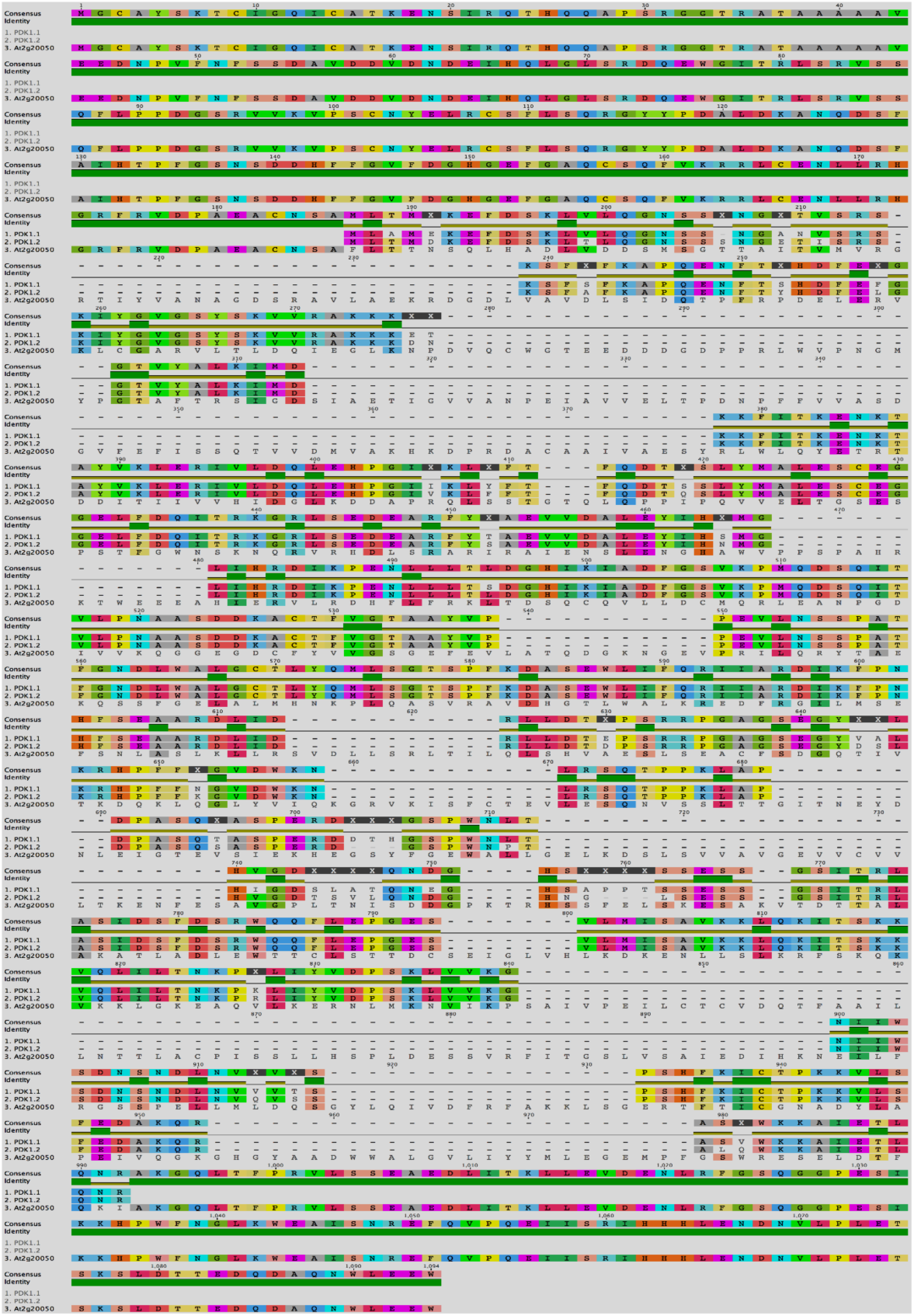
Protein alignment of PDK1.1, PDK1.2, and At2g20050. A previous study had suggested a third *PDK1*-like gene for *Arabidopsis thaliana* (At2g20050) {Zulawski et al., 2014, #17228}. However, protein sequence comparison failed to confirm any similarity of At2g20050 with *PDK1*. The amino acid alignment was performed using ClustalW algorithm and BLOSUM62 matrix in Geneious 11.1.5 software (https://www.geneious.com). Amino acids are highlighted in color. Lines represent gaps. Please note that At2g20050 consists of 1,094 amino acids (PDK1.1 491, PDK1.2486).

**S2 Fig.**
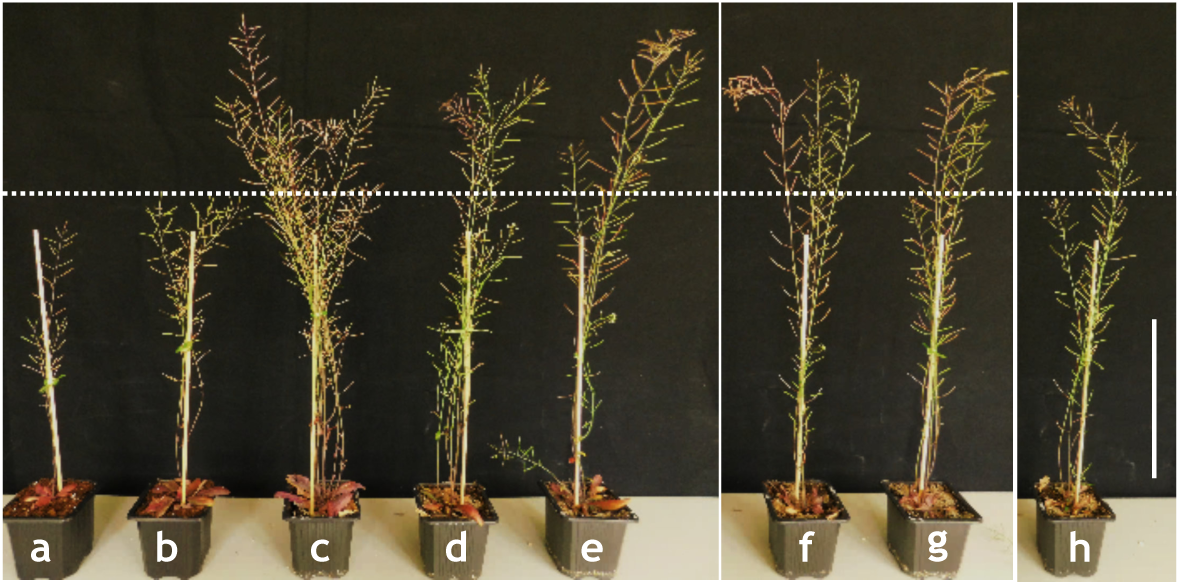
Restoration of *pdk1.1 pdk1.2* phenotype by transgenes expressing translational fusions of PDK1 to EGFP. (a) *pdk1.1-1 pdk1.2-2* (b) *pdk1.1-2 pdk1.2-3*. (c) Col-0. (d) *pdk1.1-1 pdk1.2-2 p16::gPDK1.1:EGFP*. (e) *pdk1.1-1 pdk1.2-2 p16::gPDK1.2:EGFP*. (f) *pdk1.1-2 pdk1.2-3 pUBQ10::EGFP:gPDK1.1*. (g) *pdk1.1-2 pdk1.2-3 pUBQ10::EGFP:gPDK1.2*. (h) *pdk1.1-1 pdk1.2-2 pPDK1.2::gPDK1.2:EGFP*. All constructs restore plant height. Dashed line indicates maximal plant height of the double mutants. Scale bar: 10 cm.

**S3 Fig.**
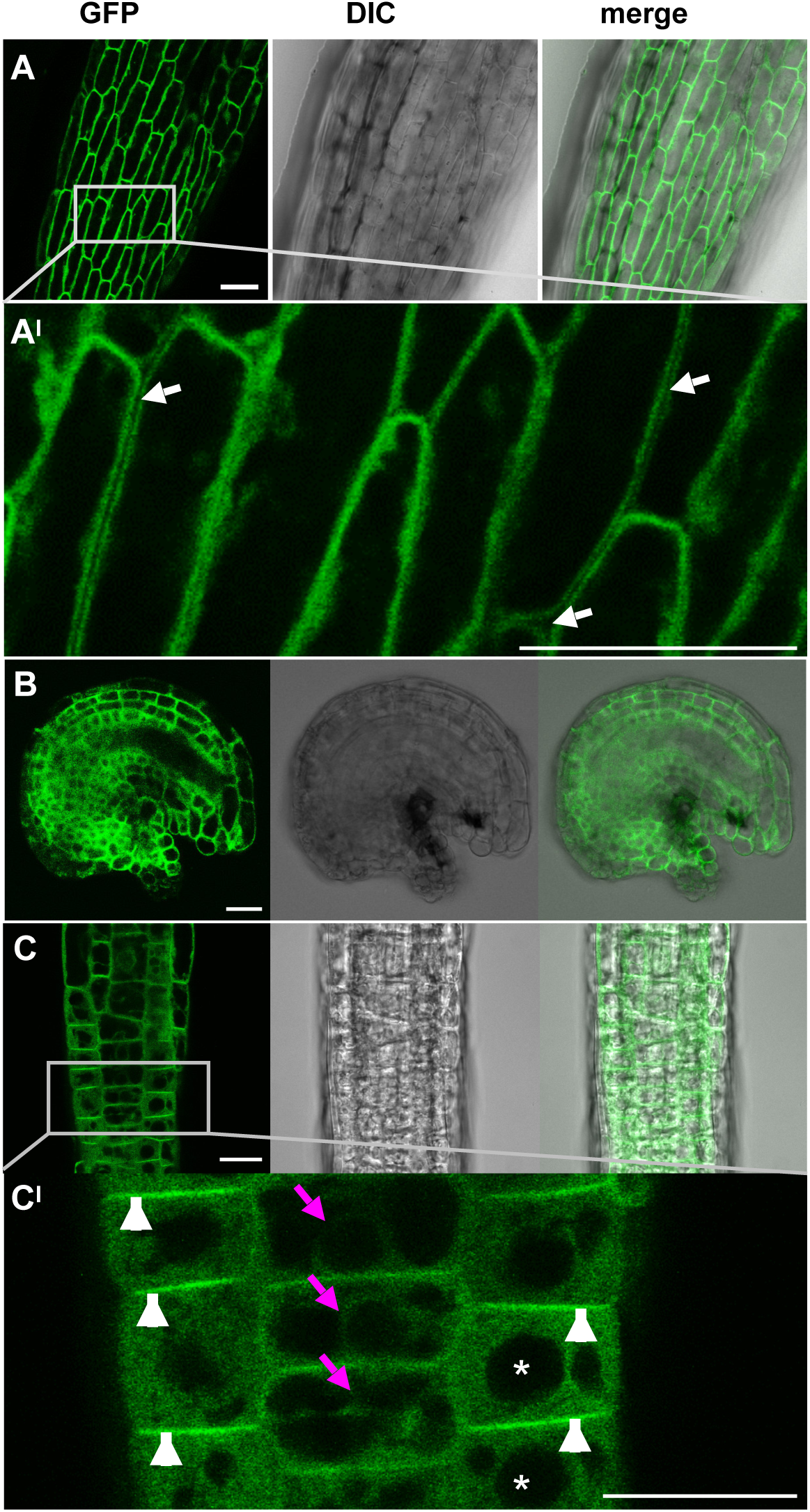
Subcellular localization of pUBQ::PDK1:EGFP reporter signals. Confocal micrographs are shown. (A) pUBQ10::PDK1.2:EGFP. (B, C) pUBQ10::PDK1.1:EGFP. (A) Filament, (A^I^) higher magnification of the region in (A), (B) ovule, (C) transition zone of a root, and (C^I^) higher magnification of the region in (C). Arrows in (A^I^) indicate the absence of GFP signal at the cell walls, magenta arrows in (C^I^) indicate most likely endoplasmic reticulum (ER), arrowheads indicate plasma membranes and asterisks indicate nuclei. Scale bars: 20 µm.

**S4 Fig.**
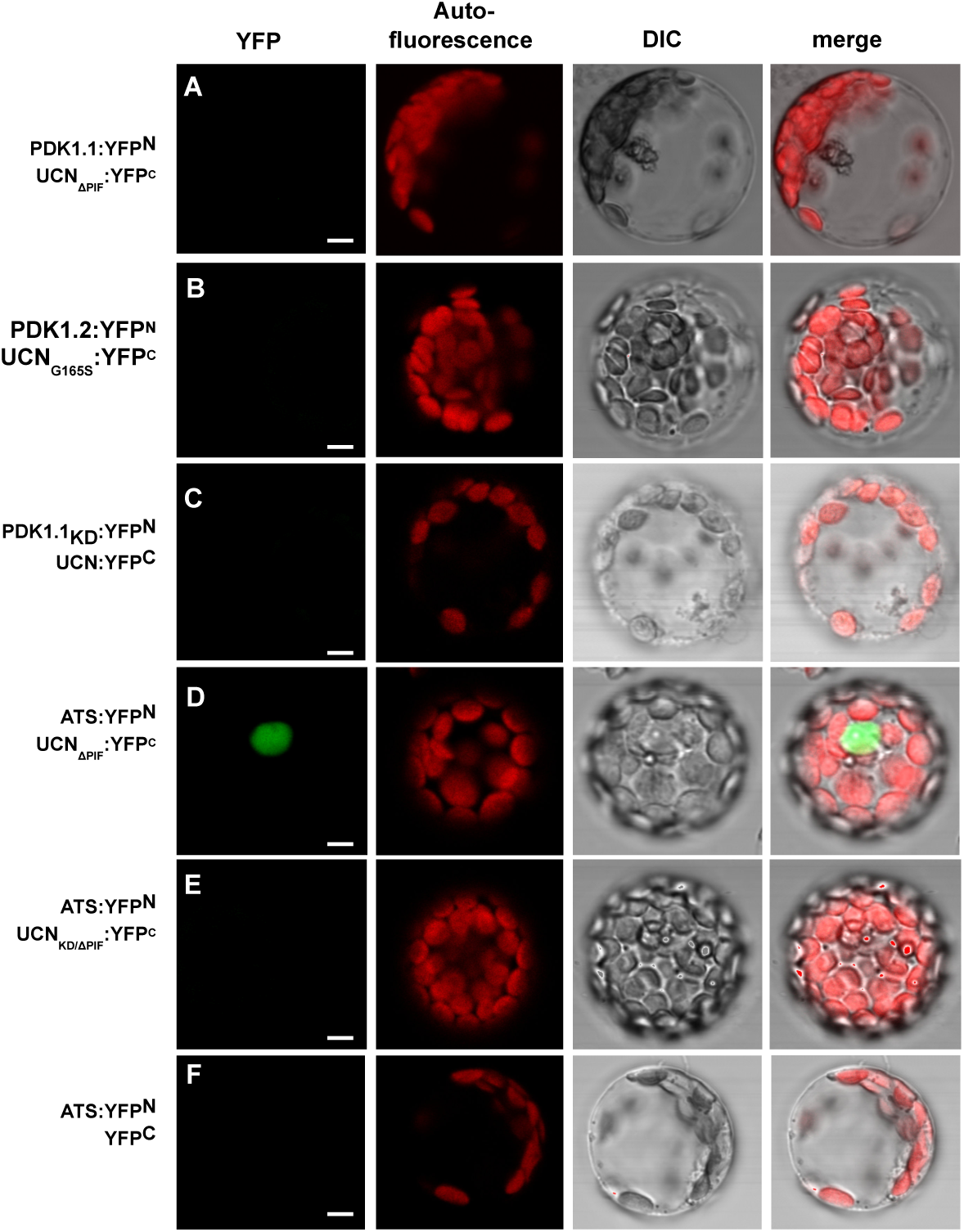
BiFC assays of PDK1 with mutant versions of UCN. Arabidopsis mesophyll protoplasts were used. The N-terminal part of YFP was fused to PDK1.1, PDK1.2, or UCN, respectively, and the C-terminal part of YFP was fused to different mutant versions of UCN. The variants are indicated. (B-C) Note absence of signal (n > 1500). (D) Signal is localized to the nucleus (126/425 scored protoplasts). Also compare to Fig. 4 in [9]. (E, F) No signal is observed (n > XX). N = 3. Scale bars: 5 µm.

**S5 Fig.**
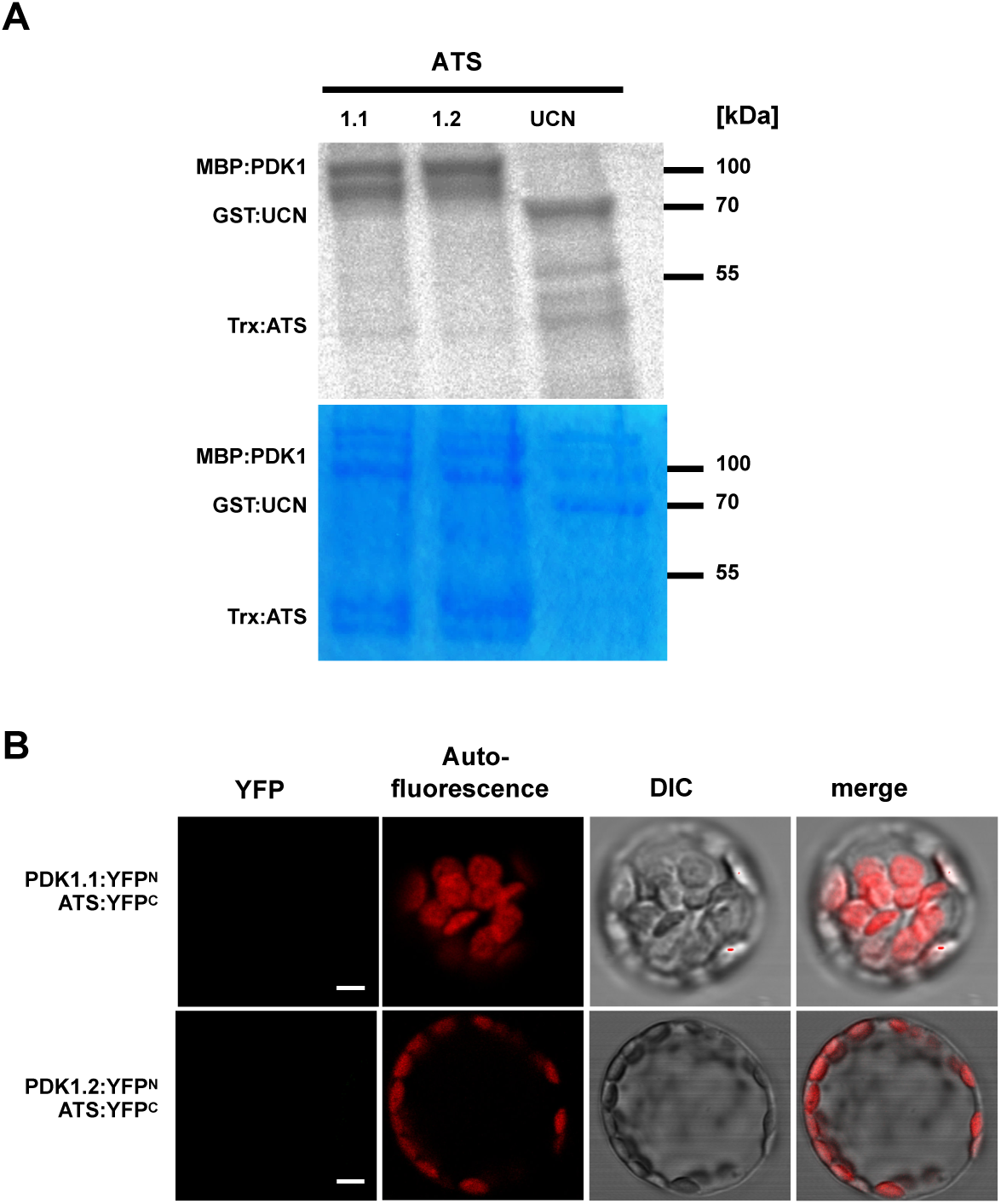
PDK1 phosphorylates ATS in vitro. (A) Kinase assays. Upper panel depicts autoradiogram. Lower panel shows corresponding coomassie brilliant blue (CBB)-stained gel. (upper panel) and autoradiograms (upper panel). Assays were performed for MBP:PDK1.1, MBP:PDK1.2, and GST:UCN, respectively, in combination with Trx:ATS. There is weak Trx:ATS phosphorylation signal in combination with MBP:PDK1 in comparison to GST:UCN/Trx:ATS (note the differences in the amount of Trx:ATS involved in the reactions). (B) BiFC assays of PDK1 and ATS. Note absence of signal. N = 3. Abbreviation: DIC, differential interference contrast. Scale bars: 5 µm.

